# Early-life stage phenomic prediction of field agronomic traits across breeding cycles in intermediate wheatgrass

**DOI:** 10.64898/2026.08.28.747871

**Authors:** Zachary N. Harris, Jackson Braley, Eric Cassetta, Jared Crain, Lee DeHaan, David Van Tassel, Allison Miller, Matthew J. Rubin

## Abstract

Perennial grains represent a promising frontier for sustainable agriculture, but breeding progress is constrained by the accessibility of genotyping and the difficulty of evaluating complex traits expressed for multiple years after establishment across heterogeneous environments. Phenomic selection may help address these challenges by using scalable, high-dimensional phenotypes collected early in development, although the robustness of such predictions across breeding cycles remains uncertain. Here, we compared genomic and phenomic selection across two breeding cycles of *Thinopyrum intermedium* (intermediate wheatgrass; IWG; Kernza®), comprising approximately 2,280 individuals from maternal half-sib families evaluated across multiple field sites and years. We constructed relationship matrices from genomic markers and early-life stage phenomic data, including seed and leaf color (HSV), CropReporter multispectral reflectance, and hyperspectral reflectance sensors. Genomic models provided the strongest predictions on average across all field traits in both cycles. Among phenomic predictors, leaf HSV was consistently the most informative, whereas CropReporter and hyperspectral data showed lower and more trait-dependent performance. Seed HSV provided little predictive value. Genomic, leaf HSV, and CropReporter models transferred across breeding cycles with little loss of predictive ability relative to within-cycle validation, demonstrating that their predictive signals were not restricted to a single breeding cycle. Despite limited similarity among relationship matrices, multi-relationship-matrix models rarely improved prediction beyond the stronger constituent single-relationship-matrix model. Together, these results show that early-life stage phenomic data provide reproducible information about agronomic performance expressed years later, but that predictor complexity and data integration do not guarantee improved prediction.

**Plain language summary:** Perennial grain crops such as intermediate wheatgrass can take several years to evaluate in the field, slowing breeding progress. We tested whether measurements collected from seeds and young seedlings could predict agronomic traits expressed years later in the field. In two breeding cycles, genomic prediction was strongest overall, but leaf color measurements from RGB images provided modest and reproducible predictions across cycles. More complex multispectral and hyperspectral measurements were generally less consistent, and combining multiple predictor data types rarely improved prediction. These results suggest that inexpensive early-life phenotyping may help breeders prioritize plants before mature field traits are available, although it is unlikely to replace genomic selection when genomic resources are already well established.

## Introduction

Developing crops that deliver high yields and ecosystem services is one of the central challenges in building a more sustainable agricultural future. Perennial herbaceous species are especially promising: their large, persistent root systems enhance carbon sequestration, stabilize soils, and reduce dependence on chemical inputs (Asbjornsen et al., 2014; Eastburn et al., 2018; Werling et al., 2014; S. Zhang et al., 2022). Yet after more than 10,000 years of attention to annual crop domestication, perennial herbaceous species remain largely undomesticated (Van Tassel et al., 2010). This lag reflects several obstacles: long juvenile phases that delay grain harvest (especially if temperate grains are sown in the spring (Jungers et al., 2022)), high genetic load, self-incompatibility, changes in shoot density during stand age, and increased risks of parasite and pathogen accumulation over multi-year lifespans, among others (Chapman et al., 2022; Smaje, 2015). From a practical breeding perspective, the need for multi-year evaluation of the same genotypes to capture age-and environment-dependent trait expression can further delay the breeding process. If perennial herbaceous crops are to help meet sustainability and production goals, improved breeding strategies that advance species through the domestication pipeline to address these barriers are essential.

The vision of an agriculture system powered largely by diverse mixtures of herbaceous perennial grains was brought to the forefront by Wes Jackson in the 1980s (W. Jackson, 1980), prompting an extensive search for candidate species that could support a more sustainable food system (Wagoner & Schaeffer, 1990). Among these, *Thinopyrum intermedium* (Host.) Barkworth & D.R. Dewey (commonly known as intermediate wheatgrass; IWG; Kernza®) emerged as a promising perennial grain candidate with large, easily threshable seeds and an extensive root system with potential for ecosystem services (Crain et al., 2024; Wagoner, 1990). Early efforts confirmed that substantial genetic gains could be achieved in IWG over a few breeding cycles, with a 77% increase in grain yield demonstrated after only two initial cycles of selection (DeHaan et al., 2014). Subsequent work has established IWG’s potential as a dual-use crop, producing grain yields alongside biomass for forage while delivering ecosystem services belowground (Culman et al., 2023; Rusch et al., 2025; Sakiroglu et al., 2020). Since initial evaluations, multiple IWG breeding programs have been developed (Bajgain et al., 2022). A subset of these programs has increasingly relied on genomic selection (GS) to reduce the length of the breeding cycle (from multiple years per cycle to two cycles per year) and to increase genetic gains (Crain, Haghighattalab, et al., 2021a; X. Zhang et al., 2016). While powerful, GS can be costly for large-scale applications that evaluate thousands of individuals each selection cycle, generating recurring expenses year after year. Approaches that can efficiently prioritize families or individuals for genotyping, or substitute recurrent genotyping with low-cost or single-time investments, are therefore especially attractive for IWG and other emerging crop breeding programs.

One promising approach is phenomic selection (PS), which adapts prediction frameworks developed for genomic selection to high-dimensional phenomic measurements rather than molecular markers (Rincent et al., 2018a). Throughout this manuscript, we use phenotype to refer to the primary agronomic traits being predicted and phenomic to refer to high-dimensional secondary phenotypes used as predictors in PS models, following terminology used in previous phenomic-prediction studies (Adunola et al., 2024; Mbebi et al., 2022). Like GS, PS can use highly parameterized linear mixed models (Endelman, 2011) or relationship-matrix and kernel-based approaches (Montesinos-López et al., 2021a; Pérez & de los Campos, 2014; X. Wang et al., 2015) to predict complex traits without requiring identification of individual causal predictors. Near-infrared and hyperspectral reflectance have been particularly common phenomic data sources because they provide rapid, high-dimensional measurements of biochemical and structural variation.

Direct comparisons between GS and PS have shown that phenomic information can sometimes approach or exceed genomic prediction, although performance depends strongly on the predictor, population, environment, and prediction design. Rincent et al. (Rincent et al., 2018b) found NIR-based predictions in wheat and poplar that were comparable to predictions from molecular markers, including across environments, while hyperspectral relationship matrices in wheat performed similarly to or better than genomic or pedigree-based models in several scenarios (Krause et al., 2019). Phenomic models have also exceeded genomic models for some growth traits in hybrid coffee (Mbebi et al., 2022), field-measured yield in wheat (R. Jackson et al., 2023), and yield in *Coffea canephora* (Adunola et al., 2023), whereas multi-environment rice analyses found PS and GS to be broadly comparable under some prediction designs (de Verdal et al., 2024). Much of this success, however, has involved phenomic measurements collected from mature plants or harvested tissues, including wheat and maize grain (Lane et al., 2020; Rincent et al., 2018c), or measurements collected relatively near the developmental period in which the target phenotype was expressed. Important exceptions demonstrate transfer across years, environments, and even generations: spectra collected several years apart from target phenotypes retained predictive value in grapevine (Brault et al., 2022), and parental NIR spectra were competitive with genomic information for predicting hybrid performance in rapeseed (Roscher-Ehrig et al., 2024). Predicting complex agronomic traits from seed or early-seedling measurements collected one or more years before trait expression nevertheless represents a substantially longer developmental and environmental separation, and more closely reflects the early-stage selection problem faced in perennial breeding.

While many phenomic selection studies to date have employed hyperspectral reflectance, other phenomic data streams have been useful in PS models. For example, germination proportion and germination timing were shown to be predictable from HSV color profiles of the seed in IWG and other perennial herbs (Keaggy et al., 2025). Before PS was officially named in 2018, techniques like metabolic-GWAS or metabolite-QTL were used to similarly connect metabolomic data with agronomic traits of interest in buckwheat (Zargar et al., 2023), wheat (Shi et al., 2020) and maize (Riedelsheimer et al., 2012). Multispectral reflectance (broad band reflectance, usually collected from canopies) and computed indices have been used to predict important traits like lodging, yield, and disease in wheat (Chandel et al., 2019; Li et al., 2020; Sharda et al., 2025) and fruit quality in strawberry (Sleper et al., 2025). Together, these studies illustrate that multiple, distinct phenomic data streams can capture biological information relevant to plant performance. However, most of these studies focus on a single phenomic data stream and on traits measured in close temporal proximity to phenotyping the traits of interest. Comparative analyses that evaluate multiple phenomic data streams side-by-side, and that ask how these streams can be combined to predict a diverse set of traits across developmental stages, years and environments, are rare, particularly in emerging perennial grains such as IWG.

In this study, we compared phenomic selection (PS) and genomic selection (GS) across two breeding cycles of intermediate wheatgrass, comprising approximately 2,280 individuals from 95-96 maternal families per cycle. Our goals were to 1) compare GS and PS within breeding cycles, 2) evaluate the transferability of both approaches across cycles, and 3) determine whether combining breeding cycles or predictor streams reliably improved predictive performance. We evaluated early-life stage phenomic data derived from seed and leaf HSV color, CropReporter multispectral reflectance and indices, and cycle-specific hyperspectral sensors under well-watered and water-limited seedling conditions. These measurements were used to predict agronomic traits expressed one to two years later across two field environments, providing a stringent test of temporally separated phenomic prediction. We partitioned major sources of variation in predictor and field traits, constructed cycle-specific phenomic relationship matrices, and compared single-and multi-relationship matrix PS models with GS. We then evaluated cross-cycle prediction and the effects of combining cycles on predictive ability. This framework allowed us to identify which early-life stage phenomic signals provided reproducible information for selection and whether additional data sources contributed complementary predictive value in an emerging perennial grain.

## Methods

### Seed sourcing and color profiling

Intermediate wheatgrass is a perennial, cool-season grass native to Eurasia that was introduced to North America in the early twentieth century for forage and erosion control. It was later selected for domestication as a grain crop because of its perennial growth habit, edible seed, and relatively large grain, with modern breeding programs now targeting yield, seed size, threshability, shatter resistance, and other domestication traits (Bajgain et al., 2022; Crain et al., 2024).

Seeds for this study were acquired from The Land Institute as part of the IWG breeding program Cycles 11 and 12 (TLI-Cycle 11-12). Seeds were acquired from random intermating of 95 and 96 known maternal families, respectively. As we reported in Keaggy et al (Keaggy et al., 2025), 2470 (Cycle 11) and 2496 (Cycle 12) seeds were scanned in a grid layout using an EPSON DS-50000 scanner (Nagano, Japan) at 1200 DPI with a SpyderCheckr® color card (datacolor, Lawrenceville, New Jersey, United States) and a ruler included. PlantCV v4.0 (Schuhl et al., 2025) was used to segment seeds from the grid layout and extract 692 features related to HSV color profiling. Seed HSV data were later subsetted to the ∼1140 seedlings per cycle used for this study.

### Germination and plant growth

For each cycle, plants were treated nearly identically, offset by a year (Cycle 11 in 2021 and Cycle 12 in 2022), at the Danforth Plant Science Center (St. Louis, MO, USA). From each family, 26 seeds were soaked in room temperature water for 24 hours before being planted in 7.6 cm wide by 20.3 cm tall square MT38 Steuwe and Sons pots (Tangent, Oregon), filled with Berger BM7 HP soil (Saint-Modeste, Quebec, Canada) supplemented with 1.5 lb/cu yd 14-14-14 Osmocote (Bloomington, Indiana) at a density of 2 seeds per pot (n = 1235 pots (Cycle 11) and n = 1248 pots (Cycle 12)). Seeds were allowed to germinate in a Conviron MTPC144 (Winnipeg, Manitoba, Canada) growth chamber in the Danforth Center Plant Growth Facility where they were bottom-watered and misted twice daily to maintain saturation (13h photoperiod, 25°C/20°C, 60% humidity, 300 µmol m ² s ¹ light). Following emergence, pots were thinned to one seedling and maintained in the growth chamber for three weeks.

Within each cycle, seedlings were split into two groups of 570 per cycle, one for each water treatment: well-watered where soil was maintained at full soil capacity and water dry-down where soil was allowed to dry and was then rewatered to prescribed set points (Supplemental Figure 1). Water treatments were imposed for 22 d in an automated Conviron GH680 growth house connected to a LemnaTec Scanalyzer 3D imaging system at the Bellwether Phenotyping Facility at the Danforth Center. The treatment periods were 16 August– 7 September 2021 for Cycle 11 and 15 August–6 September 2022 for Cycle 12. Environmental conditions matched those of the germination chamber except that light intensity was increased to 350 µmol m ² s ¹. Plants were staked to prevent interference with the automated conveyor system.

Plants automatically moved daily from the growth house via Bosch Rexroth conveyor belt system where they were imaged from the top and side with a 5 MP RGB camera. Side view images were collected in two orientations (0° and 90°). PlantCV was used to segment plant material from non-plant material and compute 692 HSV features and basic morphological summaries (e.g., plant area) for each plant daily. For each plant, for each day, we selected the HSV color profiles from the side-view orientation with maximal plant area. While this is a whole-plant above-ground phenotype, we refer to the data collected here as ‘leaf HSV’ as leaf was most of the visible area in the segmented images. We evaluated information overlap using a Mantel-like test (Mantel correlation with no permutation to assess significance (Mantel, 1967)) and phenomic selection (PS) performance across all imaging days and observed only minor differences among days (Supplemental Figure 2). To align these data with the other sensors, which were measured near the end of the treatment period, subsequent analyses used HSV features from the final three imaging days, when plants were approximately six weeks old. These observations were incorporated into the stream-specific mixed models to obtain adjusted plant-level predictor values.

### Multi-and hyperspectral reflectance

Following growth and phenotyping in the Bellwether Phenotyping Facility, plants were returned to the original growth chamber at approximately 6 weeks old. Leaf multispectral data were collected on a Phenovation CropReporter (Wageningen, The Netherlands) from one fully expanded, representative leaf detached from each plant. Multispectral reflectance was collected in the following broad bands: blue (∼470 nm), green (∼530 nm), SpcGreen (∼550 nm), red (660 nm), far red (∼730 nm), and NIR (∼790 nm). From these broadband reflectance measurements, a variety of broadband spectral indices and known physiological models were computed including anthocyanin index (AriIdx), chlorophyll index (ChlIdx), NDVI, Chl, F0, and Fv/Fm. Variance across the leaf surface for Chl and Fv/Fm were also computed. Finally, average hue, saturation, and value were reported.

Leaf hyperspectral reflectance was measured using two sensors in each breeding cycle. Cycle 11 plants were evaluated with the LinkSquare 1 (440–1000 nm; 400 bands) and LinkSquare NIR (700–1000 nm; 348 bands) sensors (Stratio, Inc., San Jose, CA, USA), whereas Cycle 12 plants were evaluated with the Innospectra NIR-S-G1 (900–1700 nm; 239 bands; Innospectra Corporation, Hsinchu, Taiwan) and SCiO Mini (740–1070 nm; 330 bands; SCiO NIR, Hod-HaSharon, Israel). For all sensors, measurements were collected from healthy, fully expanded leaves attached to the plant by placing the sensor window against the adaxial leaf surface and supporting the abaxial surface with cardboard wrapped in black fabric to standardize background reflectance. Four replicate scans were collected per plant across distinct representative leaves. LinkSquare 1, LinkSquare NIR, and Innospectra measurements were collected using an Android tablet running the Prospector application (Rife et al., 2021), with sensors connected by Bluetooth. The SCiO Mini was fitted with a rubber light shield and operated using the proprietary SCiO application.

Following data collection, spectra were analyzed using the R package waves (Hershberger et al., 2021). Outlier scans were detected by comparing the Mahalanobis distance of each reflectance curve to the total distribution of scans. Curves exceeding the 95th percentile of a chi-square distribution with degrees of freedom equal to the number of wavelengths were excluded. Remaining replicate scans were averaged within plants to produce one reflectance profile per plant and sensor. All spectral pretreatments available in waves were evaluated as candidate predictors, including untransformed reflectance (Supplemental Figure 3). Because differences in predictive performance between pretreatments were minor, we used raw reflectance values in all subsequent analyses. These features were then adjusted for cycle-and sensor-specific sources of nuisance variation.

### IWG Field Phenotyping

Plants from Cycle 11 and Cycle 12 were transplanted from the Danforth Center Plant Growth Facility to field sites near Salina, Kansas, and in the St. Louis metropolitan area, Missouri, with 0.91-m spacing between plants. Cycle 11 was transplanted on October 22, 2021, in Salina, Kansas, and September 30, 2021, in O’Fallon, Missouri. Cycle 12 was transplanted on October 6, 2022, in Salina, Kansas, and September 22, 2022, at the Danforth Center Field Research Site in St. Charles, Missouri. The O’Fallon site and the Danforth Center Field Research Site are located approximately 19 miles apart. The O’Fallon site is on Hurst silt loam soil, whereas the Danforth Center Field Research Site is on Desioux silt loam; both sites have been under agricultural production for decades, most recently in a corn-soybean rotation. The Salina, Kansas, sites were adjacent and located on McCook silt loam soil that had been used for more than 100 years for annual grain production, primarily winter wheat.

Following transplanting, plants were irrigated for 1–2 weeks, after which no supplemental irrigation was applied. Each plot was surrounded by a border row of nonexperimental intermediate wheatgrass plants. Plants were cut back in late fall or early winter each year, and fertilizer was applied after the first growing season. Areas between experimental rows were weeded or mowed throughout the growing season to reduce competition. Plants were evaluated in the field over two to three years for the traits summarized in Supplemental Table 1, following the trait-specific methods described by (Crain, DeHaan, et al., 2021a; Crain et al., 2022; Crain, Haghighattalab, et al., 2021b).

### Variance partitioning and adjustment of field and phenomic traits

For each field trait measured we sought to explore variation across maternal family, adult environments (KS vs STL; site), and sample years (where available) within each breeding cycle. Linear models were fit to each scaled (z-score) field trait with family, site, and year as fixed interacting effects. From each model we calculated the variance explained by each term as η^2^ = SS_term_ / SS_total_ (multiplied by 100 to compose a percentage). A separate watering-treatment-only model was used to affirm that there was no residual watering treatment effect in the field trait expression. A best linear unbiased prediction (BLUP) was estimated for each individual using ASREML (Gilmour et al., 2015). The mixed model accounted for random effects of site, year, maternal influence, and individual genotype. Additionally, a two-dimensional autoregressive (AR1xAR1) was fit to model spatial variation between rows and columns, following standard practice in IWG breeding trials to derive stable trait estimates across years and environments (Crain et al., 2020, 2022). This approach produces shrunken predictions that preserve genetic signal while absorbing environmental, spatial, and experimental variation across years and locations.

We applied a parallel variance-partitioning framework to the unadjusted phenomic predictor features, including each HSV feature, each CropReporter reflectance band and broadband index, and reflectance at each wavelength from all hyperspectral sensors. Because these measurements were collected before field establishment, watering treatment replaced field site in the predictor models. Additional terms were included as appropriate to quantify variation associated with technical and procedural factors, such as sampling date, image-collection day, plate position (CropReporter), and device identity (hyperspectral sensors). Variance explained by each term was calculated as described above. To provide a higher-level summary of variation within each phenomic data stream, we also performed principal component analysis and applied the same variance-partitioning framework to all resulting principal components. For each model term, the overall percent variance explained was calculated as the variance-weighted mean across principal components, using the proportion of total predictor variance captured by each component as the weight.

For within-cycle analyses, we estimated adjusted predictor values for each feature with nonzero variance within each phenomic data stream using sommer (Covarrubias-Pazaran, 2016). Models accounted for plant ID and maternal family as random effects as well as other stream-specific nuisance sources of variation. Adjusted values were obtained as predicted values for each individual plant, thereby removing nuisance effects while retaining both family-level and within-family biological variation. Interaction terms were included only when the model design was sufficiently balanced across the corresponding factor combinations. The adjustment model differed among data streams to reflect their respective collection protocols. Leaf HSV features were adjusted for watering treatment and image-collection day. CropReporter reflectance and broadband indices were adjusted for watering treatment, sampling date, sampling plate, position within plate, family-by-treatment interactions, and family-by-sampling-date interactions. LinkSquare, LinkSquare NIR, and SCiO Mini features were adjusted for watering treatment, scan date, and device identity. Innospectra features were adjusted for the same effects, together with interactions between maternal family and watering treatment, scan date, and device identity.

In cross-cycle and combined-cycle models both field traits and predictor data streams (seeds HSV, leaf HSV, CropReporter) were additionally adjusted for cycle as a random effect.

### Phenomic selection modeling

Phenomic selection models were fit following the methods of Krause et al (Krause et al., 2019) and Keaggy et al (Keaggy et al., 2025). Within each breeding cycle, phenomic relationship matrices were constructed separately for each adjusted data stream using a linear kernel standardized by the number of features: tcrossprod(X) / ncol(X), where X was a centered and scaled matrix of adjusted phenomic features. Separate matrices were generated for each data stream, including seed HSV profiles, leaf HSV profiles, leaf CropReporter traits, leaf and hyperspectral reflectance. Because the hyperspectral instruments differed between breeding cycles, sensor-derived relationship matrices were treated as cycle-specific data streams rather than pooled across instruments. PS models were fit in R using the BGLR package (Pérez & de los Campos, 2014) with each relationship matrix defining the prior covariance structure of a reproducing kernel Hilbert space term. Within-cycle predictive performance was evaluated using five-fold cross-validation. In each fold, field-trait values for 20% of individuals were set to missing and predicted from models trained on the remaining 80%. For models incorporating multiple data streams, genomic and phenomic relationship matrices, or pairs of phenomic relationship matrices, were entered as separate terms in the BGLR ETA list. Cross-cycle models were trained in one breeding cycle and evaluated in the other, whereas combined-cycle models used individuals from both cycles in the training population. Model performance was evaluated as the Pearson correlation between observed and predicted field-trait values for the relevant validation set.

### IWG Genotyping

All IWG plants were profiled using a two-enzyme genotyping-by-sequencing protocol (Poland et al., 2012; Sthapit et al., 2025). The 2280 plants were genotyped in the same manner as IWG plants in The Land Institute Breeding program which consists of collecting ∼50 mg of fresh tissue followed by DNA extraction using MagMAX Plant DNA Isolation kit (ThermoFisher Scientific) following manufacturer protocol. Genotyping-by-sequencing libraries were constructed using *PstI* and *MspI* restriction enzymes combining 192 samples into each sequence library. All libraries were sequenced at Psomagen Inc. (Rockville, Maryland), with each library (n = 6) being sequenced on a single lane of an Illumina HiSeqX machine.

Bioinformatic processing followed The Land Institute’s breeding pipeline for consistency (Crain, DeHaan, et al., 2021b). This included identifying single nucleotide polymorphisms (SNPs) using the TASSEL GBSv2 pipeline (Glaubitz et al., 2014) with the IWG draft genome version 3.1 (https://phytozome-next.jgi.doe.gov/info/Tintermedium_v3_1). Sequence reads were required to be at least 50bp long and align to only one unique location within the genome to be used for SNP calling. To call heterozygous loci, a read depth of two contrasting tags was required. We required a minimum read depth of four to call a homozygous locus. Potential loci were filtered for a minor allele frequency greater than 0.05 and less than 70% missing data. Individual plants were removed if they had more than 95% missing loci. The final data set of 114,851 markers and 2,280 individuals was imputed using Beagle version 4.1 (Browning & Browning, 2016) using default parameters. Sequence data has been uploaded to NCBI Sequence Read Archive (SRA) (https://www.ncbi.nlm.nih.gov/bioproject/) as part of BioProject PRJNA1044453.

### Genomic selection modeling

Genomic selection models were fit in BGLR using a genomic relationship matrix constructed with the A.mat function in rrBLUP (Endelman, 2011). The genomic relationship matrix was included as a reproducing kernel Hilbert space (RKHS) term defining the prior covariance among individuals. In models combining genomic and phenomic information, the genomic and phenomic relationship matrices were included as separate kernels in the BGLR ETA list.

## Results

### Genetic, environmental, and technical factors structure early-life stage phenomic data across breeding cycles

Early-life stage phenomic data showed broadly consistent patterns of genetic (family), environmental (watering treatment), temporal (date) and technical (device id) variation across the two breeding cycles (Figure 1). Maternal family effects were similar between cycles within each shared data stream. Seed HSV was dominated by maternal-family-associated variation, which accounted for 88.7% of total predictor variance in Cycle 11 and 87.3% in Cycle 12. Maternal family explained a smaller but consistent proportion of variation in leaf HSV and CropReporter data, accounting for 9.2 - 10.3% and 12.9 - 14.7% of total variance, respectively. Family-by-watering-treatment interactions contributed an additional 5.1 - 5.8% of variation in leaf HSV and 7.5 - 8.3% in CropReporter traits, indicating that a small portion of early-life stage phenotypic variation reflected family-specific responses to seedling water conditions. Watering treatment alone explained relatively little variation in any data stream or cycle (0.05 - 2.57%). Hyperspectral reflectance exhibited a similar maternal-family component, with family explaining approximately 8–11% of total variance across sensors and cycles. In contrast to the image-derived data, however, hyperspectral measurements were strongly structured by technical sources of variation. Device identity explained 46.9% and 76.3% of total variation in the Cycle 11 LinkSquare 1 and LinkSquare NIR data, respectively, and 38.1% and 24.9% in the Cycle 12 Innospectra and SCiO Mini data. These substantial technical effects motivated the stream-specific adjustment models used before construction of the phenomic relationship matrices. Additional modeled technical and procedural effects also accounted for substantial variation in several hyperspectral streams. Residual variation was consequently much lower for the hyperspectral sensors than for leaf HSV and CropReporter data, for which residuals accounted for approximately 62–75% of total variance.

**Figure 1.**
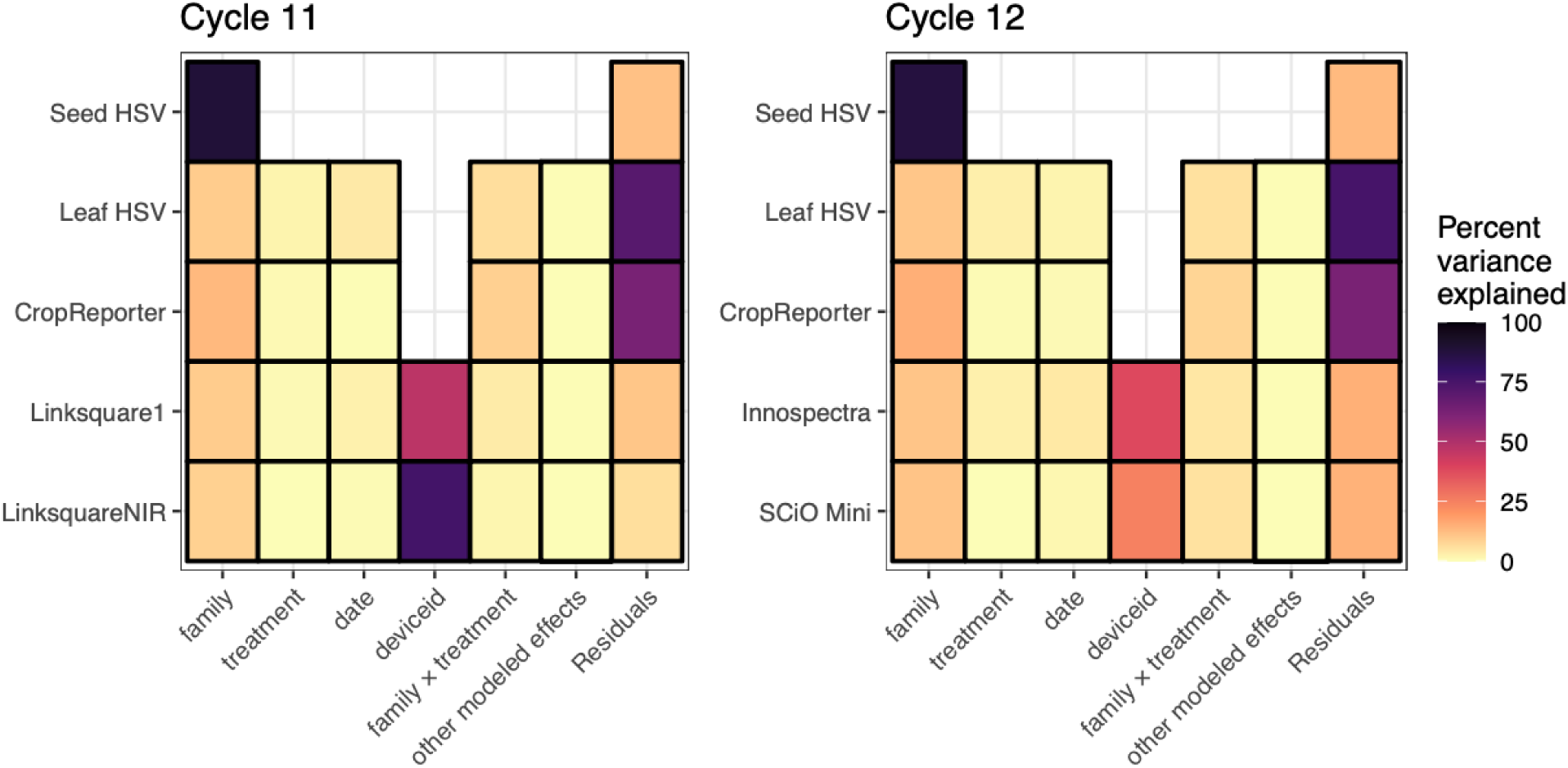
Variance partitioning of early-life stage phenomic data across two intermediate wheatgrass breeding cycles. Heatmap cells show the percent variance explained by maternal family, watering treatment, sampling or imaging date, device identity, family-by-treatment interactions, all other known and modeled effects, and residual variation within each phenomic data stream. Separate analyses were performed for Cycle 11 and Cycle 12. For each data stream, principal component analysis was performed on the unadjusted predictor features, and the same variance-partitioning model was fit to every resulting principal component within a data stream. The data stream-level percent variance explained for each model term was calculated as the variance-weighted mean across all principal components, using the proportion of total predictor variance captured by each component as its weight. Seed HSV models included only maternal family because seed color was measured before seedlings were assigned to experimental treatments. “Other modeled effects” combines stream-specific technical and procedural terms, including plate, plate position, and interactions involving maternal family, sampling date, or device identity; these effects are decomposed in Supplemental Figure 4. Blank cells indicate terms that were not included in the corresponding model.

Feature-level variance partitioning and decomposition of family, watering-treatment, and technical effects are provided in Supplemental Figures 4–6. The alternative watering treatments were included to test whether some environments might broaden phenotypic variation and thereby increase the information available for prediction. Treatment effects were generally localized rather than pervasive, with the clearest differences occurring in leaf HSV saturation features (Supplemental Figure 4). Feature-wise Brown-Forsythe tests identified treatment-associated differences in variance for subsets of leaf HSV and hyperspectral features, although differences were also detected in seed HSV, which was measured before treatment assignment (Supplemental Table 2). When feature variances were summarized across entire data streams, consistent treatment differences were limited to the Cycle 12 hyperspectral sensors, but the direction of change differed between instruments (Supplemental Figure 5). Likewise, watering treatment did not produce a consistent shift in the proportion of phenotypic variance attributable to maternal family (Supplemental Figure 6). Together, these analyses provide little evidence that different seedling environments broadly increased phenotypic variance or family-associated signal across predictor streams, supporting its treatment as an experimental effect to be adjusted before phenomic prediction. Consistent with the above findings, we found few significant differences between the mean phenomic selection accuracies across seedling watering treatments when evaluating each treatment independently, though these effects were not directionally consistent or practically meaningful (Supplemental Figure 7).

### Field traits exhibit reproducible family effects but substantial environmental and temporal variation across breeding cycles

Field traits showed substantial variation attributable to maternal family, field site, evaluation year, and their interactions within both breeding cycles (Figure 2; Supplemental Table 3). Maternal family significantly influenced nearly all traits, although the magnitude of family-associated variation differed among traits and between cycles. Family effects were strongest and most consistent for seed morphology traits, including seed circularity, seed length, and naked seed area. For example, maternal family explained 20.0% and 19.3% of variation in seed circularity in Cycles 11 and 12, respectively, and 19.4% and 18.2% of variation in seed length. More moderate family effects were observed for many yield, developmental, and architectural traits, generally accounting for approximately 5–18% of total variation.

**Figure 2.**
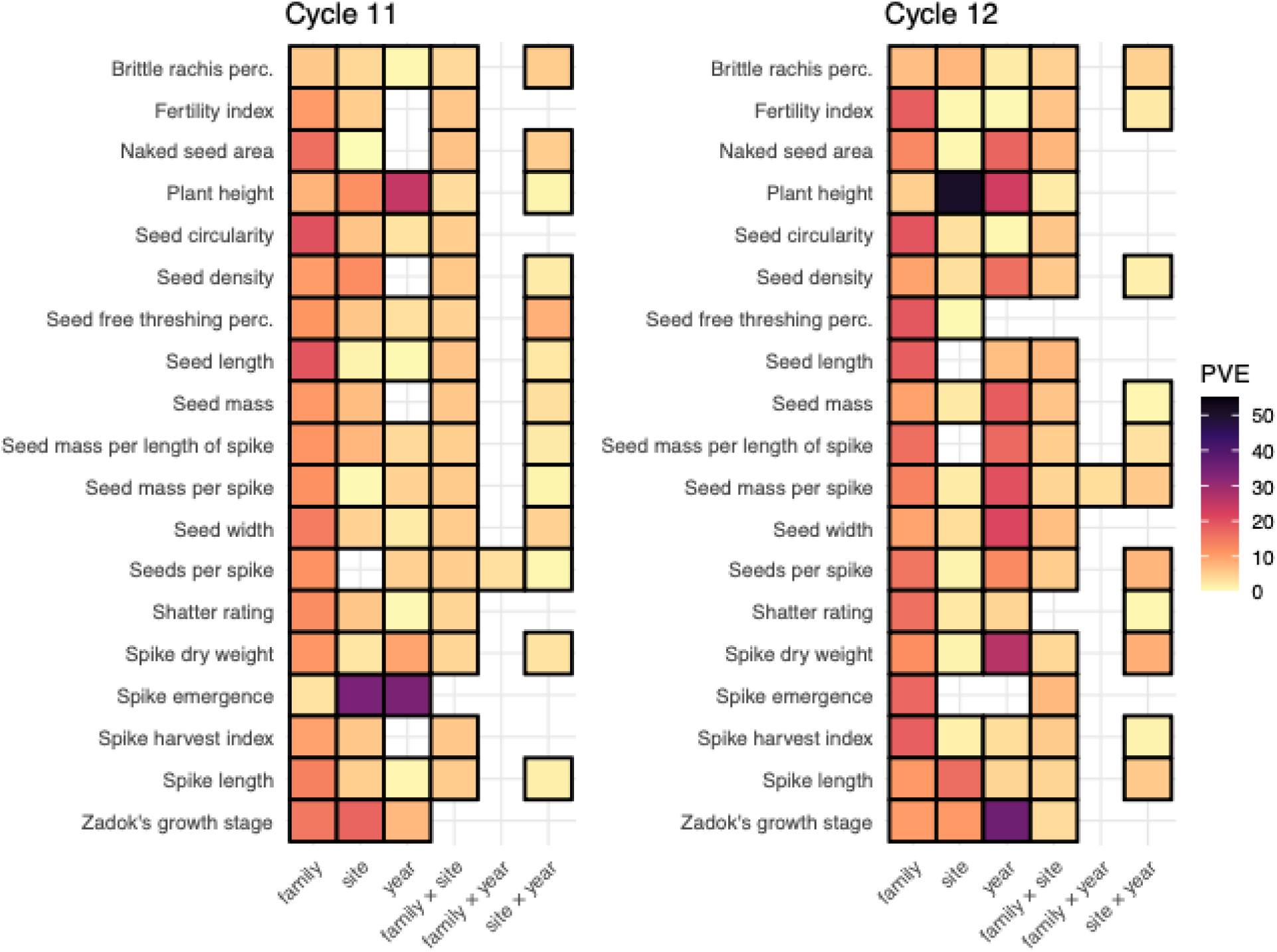
Variance partitioning of field traits across two intermediate wheatgrass breeding cycles. Heatmap cells show the percent variance explained (η^2^) by maternal family, field site, evaluation year, family-by-site interactions, family-by-year interactions, and site-by-year interactions for each field trait analyzed within Cycle 11 and Cycle 12. Field traits were analyzed directly using scaled observations, and the proportion of total variance explained by each model term was calculated as = SS_term_ / SS_total_. Blank cells indicate either that a term was not included in the corresponding model or that it was not statistically significant. One significant three-way interaction (Cycle 11 brittle rachis percentage; p = 0.02, PVE = 3.15%) was removed for visualization purposes.

Environmental and temporal effects on field traits were often larger than maternal-family effects but were less consistent across traits and cycles. Site strongly influenced spike emergence, plant height, spike length, seed density, and Zadok growth stage in at least one cycle. The most pronounced site effect was observed for plant height in Cycle 12, for which site explained 50.9% of total variation, compared with 11.5% in Cycle 11. Year effects were also substantial for several developmental and yield-related traits. In Cycle 11, year explained 34.1% of variation in spike emergence and 24.8% in plant height, whereas in Cycle 12 it explained 35.8% of variation in Zadok growth stage, 26.3% in spike dry weight, and approximately 16– 22% in several seed-size and yield traits.

Family-by-site interactions were detected for most traits in both cycles but were generally smaller than the family, site, or year effects. These interactions commonly explained approximately 3–7% of total variation, indicating that relative family performance varied across field environments. Site-by-year interactions were also present for several traits, particularly yield and reproductive traits, whereas family-by-year interactions were uncommon. Together, these patterns indicate that field-trait expression was shaped by a persistent maternal-family component superimposed on substantial and trait-specific environmental and temporal variation.

We separately tested each field trait within each breeding cycle for any residual effect of the earlier seedling watering treatment (Supplemental Figure 8); after Benjamini–Hochberg correction, no trait retained a significant treatment effect (median PVE = 2.87%). We therefore generated individual-level BLUPs, accounting for variance among sites, spatial effects within sites, years, and maternal families. We then used these environmentally adjusted phenotypes as response variables in subsequent genomic and phenomic prediction analyses.

### Genomic and leaf HSV relationship matrices are consistently the strongest predictors across breeding cycles

Across all evaluated field traits, the relative performance of genomic and phenomic predictors was broadly consistent between breeding cycles (Figure 3; Supplemental Figure 9; Supplemental Table 4). Genomic selection provided the strongest overall predictive performance, with mean predictive abilities of r = 0.31 in Cycle 11 and r = 0.29 in Cycle 12. Among the phenomic data streams, leaf HSV was consistently the strongest predictor, with mean predictive abilities of r = 0.16 and r = 0.17, respectively. CropReporter and the best-performing hyperspectral sensor in each cycle showed lower but similar mean performance: r = 0.09 for CropReporter and r = 0.09 for LinkSquare 1 in Cycle 11, and approximately r = 0.08 for both CropReporter and Innospectra in Cycle 12. LinkSquare NIR and SCiO Mini performed more poorly, with mean predictive abilities of r = 0.05 and r = 0.03, respectively, while seed HSV provided effectively no predictive ability in either cycle.

**Figure 3.**
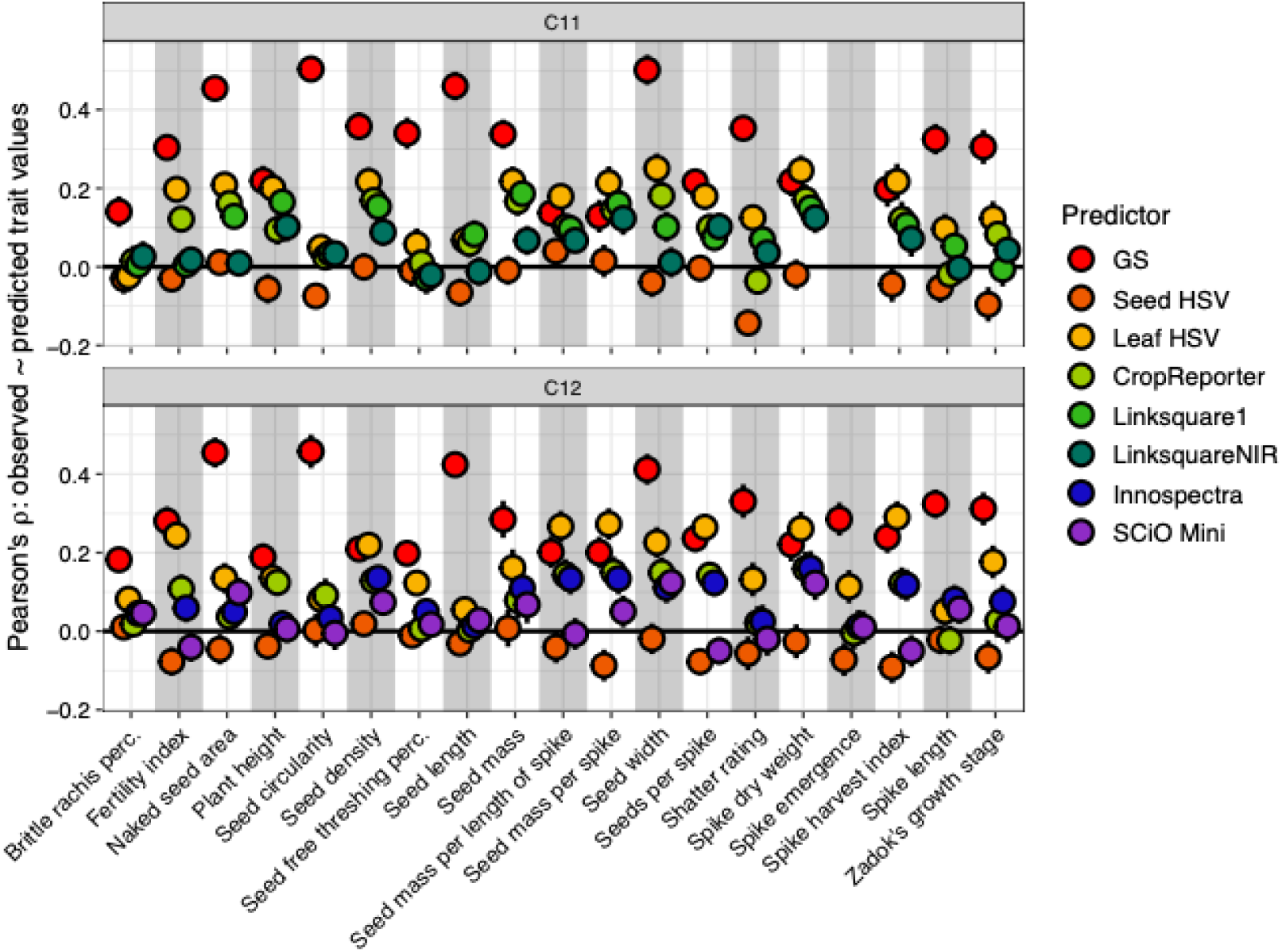
Predictive ability of genomic and phenomic selection models across field traits in two intermediate wheatgrass breeding cycles. Pearson correlations between observed and predicted trait values are shown for genomic selection (GS) and each phenomic data stream within Cycle 11 and Cycle 12. Phenomic predictors included seed HSV, leaf HSV, CropReporter traits, and cycle-specific hyperspectral reflectance data from LinkSquare 1 and LinkSquare NIR in Cycle 11 and Innospectra and SCiO Mini in Cycle 12. Points represent mean predictive ability across five cross-validation folds, and error bars denote 83% confidence intervals. Values above zero indicate positive correspondence between observed and predicted trait values.

Predictive performance nevertheless varied substantially among traits. Genomic selection performed especially well for seed shape traits, including naked seed area, seed circularity, seed length, and seed width, for which predictive ability commonly exceeded r = 0.40 in both cycles. Leaf HSV performance approached or exceeded genomic prediction for seed mass per unit spike length, seed density, seeds per spike, spike dry weight, spike harvest index, and seed mass per spike in one or both cycles. In contrast, leaf HSV was less informative for seed shape traits, for which GS retained a clear advantage.

The remaining phenomic streams showed more modest and trait-specific predictive value. CropReporter and hyperspectral relationship matrices produced positive predictions for several yield and growth traits, but their performance was generally below that of leaf HSV. Seed HSV rarely provided useful prediction of mature field traits despite its strong maternal-family structure. Overall, the same broad pattern emerged in both breeding cycles: genomic relationships were the strongest predictors of mature field performance, leaf HSV provided the most informative phenomic representation, and the remaining phenomic streams offered weaker and more variable predictive signals.

### Combining relationship matrices provides little additional predictive gain

Despite limited similarity among the individual-level relationship structures captured by genomic and phenomic data streams (Supplemental Figure 9), combining relationship matrices rarely improved prediction beyond the best-performing single relationship matrix. Across both breeding cycles, two-relationship-matrix models that combined GS with phenomic information or paired two phenomic relationship matrices generally produced predictive abilities comparable to, but rarely greater than, the stronger constituent model (Figures 3–4). In particular, although models combining GS and leaf HSV sometimes exceeded GS alone, their performance was generally indistinguishable from the better of GS or leaf HSV considered separately. Results from all evaluated multi-relationship-matrix models, including phenomic-only combinations and models containing up to all six relationship matrices per breeding cycle, are provided in Supplemental Table 5. These results provide little evidence that the evaluated relationship matrices contained complementary predictive information sufficient to improve performance through relationship matrix combination. Thus, differences in relationship-matrix structure did not necessarily translate into complementary trait-relevant signal.

**Figure 4.**
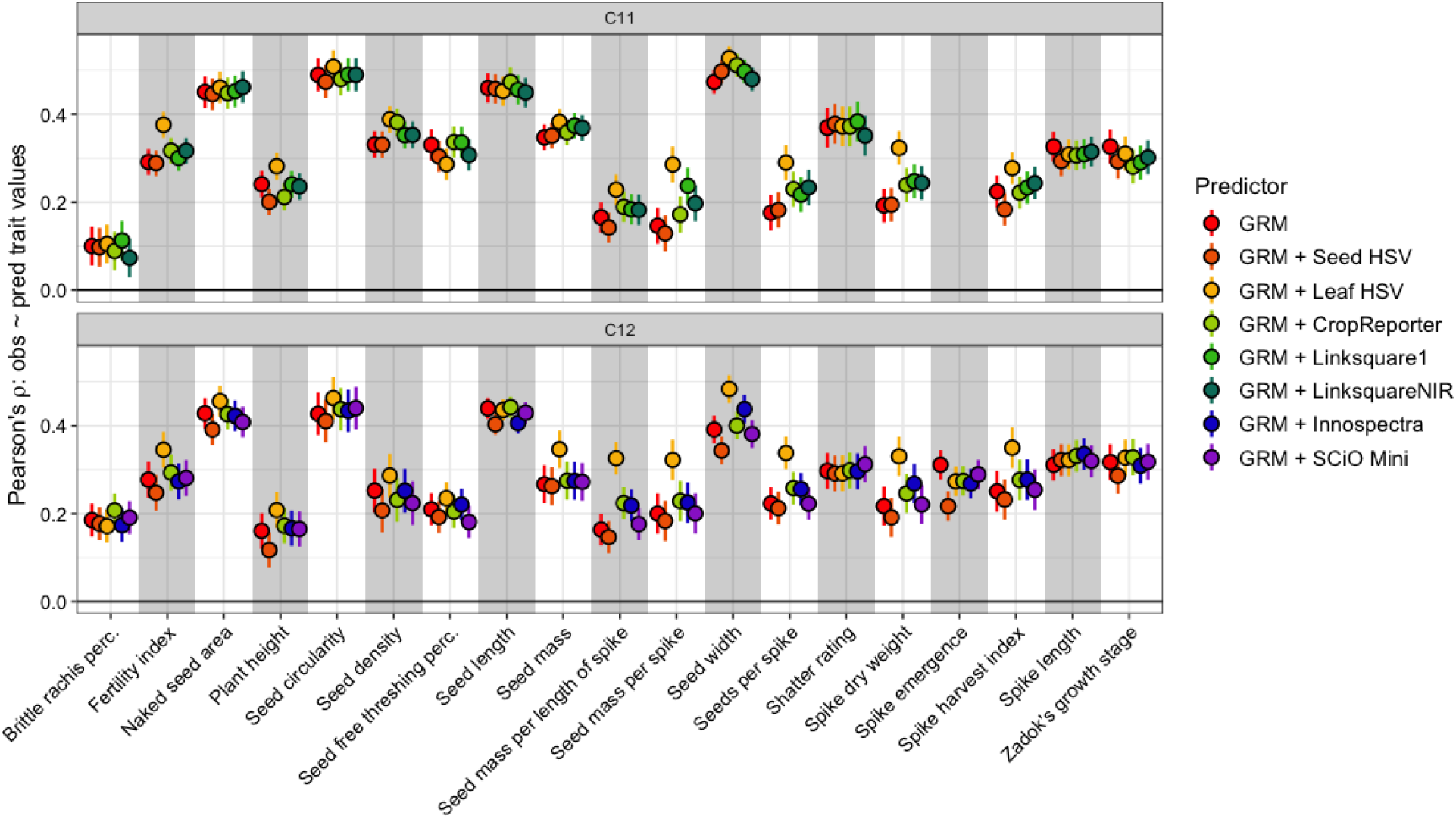
Predictive ability of genomic and combined genomic–phenomic relationship models across two intermediate wheatgrass breeding cycles. Pearson correlations between observed and predicted trait values are shown for models containing the genomic relationship matrix alone (GRM) or the genomic relationship matrix combined with one phenomic relationship matrix. Phenomic predictors included seed HSV, leaf HSV, CropReporter traits, and cycle-specific hyperspectral reflectance data from LinkSquare 1 and LinkSquare NIR in Cycle 11 and Innospectra and SCiO Mini in Cycle 12. Points represent mean predictive ability across five cross-validation folds, and error bars denote 83% confidence intervals. Models are shown for all field traits evaluated within each breeding cycle.

### Genomic and leaf HSV models transfer across breeding cycles, whereas gains from combining cycles depend on the predictor stream

Genomic prediction transferred effectively between breeding cycles (Figure 5; Supplemental Table 6). Mean predictive ability was r = 0.31 within Cycle 11 and r = 0.29 within Cycle 12, compared with r = 0.29 when models trained in Cycle 11 were used to predict Cycle 12 and r = 0.30 in the reciprocal direction. Cross-cycle performance therefore showed little loss relative to prediction within the target cycle. Trait-level performance was also highly consistent between within-cycle and cross-cycle models, with genomic prediction remaining strongest for seed shape traits such as seed circularity, seed length, naked seed area, and seed width.

**Figure 5.**
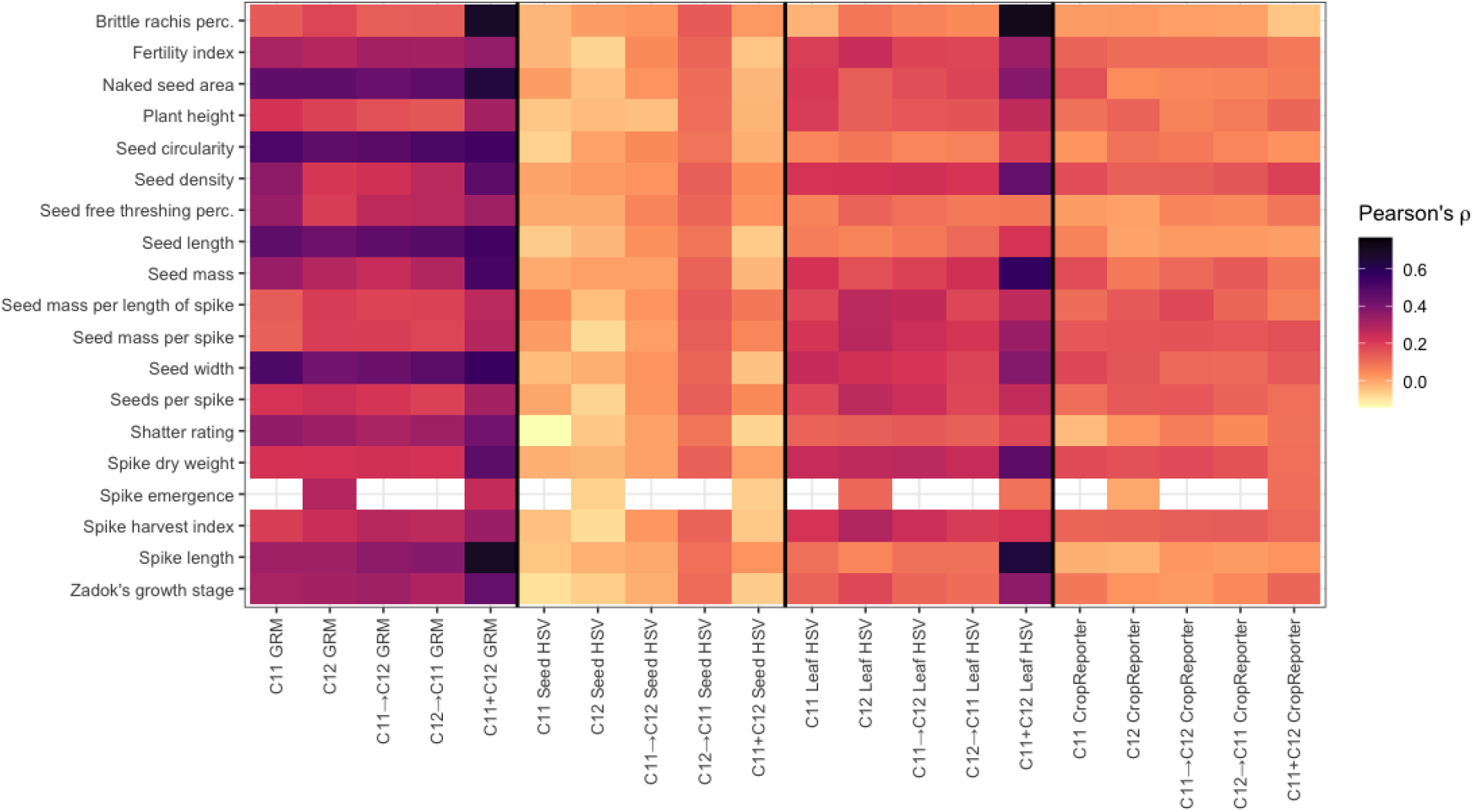
Within-cycle, cross-cycle, and combined-cycle predictive ability of genomic and phenomic relationship models across field traits. Heatmap cells show Pearson correlations between observed and predicted trait values for models based on the genomic relationship matrix (GRM), seed HSV, leaf HSV, and CropReporter data. Within-cycle models were trained and evaluated within Cycle 11 or Cycle 12 using cross-validation. Cross-cycle models were trained in one breeding cycle and evaluated in the other, with arrows indicating the direction of prediction. Combined-cycle models used data from both breeding cycles. Darker cells indicate stronger positive predictive ability, whereas values near zero indicate little correspondence between observed and predicted trait values. Blank cells indicate trait–analysis combinations that were not evaluated. Fold-level estimates and complete numerical results are provided in Supplemental Table 6.

Leaf HSV showed similarly robust transfer across cycles (Figure 5; Supplemental Table 6). Mean predictive ability was r = 0.16 within Cycle 11 and r = 0.17 within Cycle 12, compared with r = 0.17 for Cycle 11 models projected into Cycle 12 and r = 0.15 for Cycle 12 models projected into Cycle 11. The traits best predicted by leaf HSV were also broadly conserved across validation schemes, with the strongest performance generally observed for spike dry weight, seed mass, seed density, seed mass per spike, and related yield components. CropReporter models likewise transferred without substantial loss: cross-cycle predictive abilities (r = 0.090 and r = 0.09) were similar to their within-cycle values (r = 0.09 and r = 0.08). In contrast, seed HSV provided little predictive value within either cycle or when Cycle 11 was used to predict Cycle 12. The apparently positive performance of Cycle 12 seed HSV models projected into Cycle 11 was inconsistent with the other validation schemes and was therefore not interpreted as evidence of robust cross-cycle prediction.

Combining both breeding cycles increased mean genomic predictive ability to r = 0.44, exceeding within-cycle performance for nearly all shared traits. Leaf HSV also showed a substantial increase in the combined-cycle analysis, reaching a mean predictive ability of r = 0.34. However, unlike genomic prediction, the magnitude and trait distribution of this increase were inconsistent with the cross-cycle results and likely reflected residual differences between breeding cycles that could not be fully removed during predictor adjustment (Supplemental Figure 10). The combined-cycle improvement for leaf HSV should therefore be interpreted cautiously rather than attributed solely to the larger training population. CropReporter showed no corresponding benefit from combining cycles, with mean predictive ability remaining near its within-and cross-cycle values r = 0.08. Seed HSV also remained effectively non-predictive in the combined analysis. Together, these results indicate that genomic and leaf HSV relationship matrices capture signals that transfer across breeding cycles, but increasing training-population size does not uniformly improve phenomic prediction and may amplify unresolved cycle structure in some data streams.

## Discussion

Breeding perennial crops can be accelerated by making selection decisions before mature agronomic traits can be measured directly (Crain, Haghighattalab, et al., 2021a). Early-stage selection is especially challenging because agriculturally important traits (e.g., flowering and yield) are expressed only after field establishment and vary across sites and years. Similar to genomic selection, which enables early selection using genome-wide marker information, phenomic selection may provide an additional means of informing these decisions. Its usefulness in this context depends on whether phenotypes collected during seedling development contain information about mature performance and whether that information remains relevant across breeding cycles. In this study, genomic selection provided the strongest overall predictions across both breeding cycles, but the relative performance of phenomic data streams was broadly reproducible. Leaf HSV consistently provided the strongest phenomic predictions, CropReporter data showed more modest predictive ability, and seed HSV provided little useful signal despite strong maternal-family structure. Genomic, leaf HSV, and CropReporter models transferred across breeding cycles with little loss of predictive performance relative to within-cycle validation, indicating that their predictive signals were not confined to a single breeding population. In contrast, combining relationship matrices rarely improved prediction beyond the stronger constituent model, and the apparent gain from combining cycles for leaf HSV should be interpreted cautiously because residual cycle-associated structure may have contributed to its performance. Together, these results show that early-life stage phenomic data can provide reproducible information about agronomic performance expressed years later, but that neither greater predictor complexity nor the addition of more data streams necessarily yields stronger models.

### Genomic prediction outperforms phenomic prediction and generalizes across breeding cycles

Across our multi-year, multi-environment IWG trial, genomic selection provided the strongest overall predictions in both breeding cycles. This result is consistent with previous work showing that many of the traits evaluated here are moderately heritable, associated with identified QTL, and predictable using genomic selection in IWG (Altendorf et al., 2022; Bajgain et al., 2024; Crain, DeHaan, et al., 2021b; Crain et al., 2020, 2022). Genomic markers encode additive relationships that remain stable across developmental stages and environments, whereas phenomic measurements collected at a single early-life stage may become less informative as later phenotypes are shaped by development, plasticity, and genotype-by-environment interactions (Fritsche-Neto et al., 2025; Johansen et al., 2025). Genotyping also provides a persistent source of information that can be reused across years and environments within a breeding population, reinforcing the practical value of GS for complex agronomic traits expressed after establishment in perennial crops (Esfandyari et al., 2020; Fritsche-Neto et al., 2025; Seyum et al., 2022).

Genomic models trained in one breeding cycle predicted the other cycle with little loss of performance relative to within-cycle cross-validation. This transferability likely reflects the close relationship between Cycle 11 and Cycle 12, which are successive recurrent-selection cycles drawn from a breeding population with a narrow founder base and genetic variation largely constrained within the breeding program (Crain et al., 2020, 2024; X. Zhang et al., 2016). Under these conditions, marker-derived relationships and marker–trait associations established in one cycle would be expected to remain informative in the next. From an applied perspective, this result suggests that genomic training data accumulated in one cycle can retain value in subsequent cycles rather than requiring models to be rebuilt entirely from contemporaneous phenotypes. Combining cycles further improved genomic prediction, consistent with evidence that increasing the size and diversity of a genetically relevant training population can improve predictive accuracy (Edwards et al., 2019).

Direct comparisons between GS and PS have produced mixed outcomes. Phenomic prediction has equaled or exceeded genomic prediction for some growth and yield traits in coffee (Adunola et al., 2023; Mbebi et al., 2022), wheat (Winn et al., 2023), and soybean (Van der Laan et al., 2025), while genomic prediction has been clearly superior for other traits and systems, including apple (Jung et al., 2025) and soybean seed composition (Van der Laan et al., 2025). However, differences in predictive ability between PS and GS should not be interpreted directly as differences in selection accuracy or expected genetic gain, because phenomic predictions may capture both genetic and non-genetic components of trait variation and the two approaches can affect multiple components of the breeder’s equation (F. Wang et al., 2025). Within that limitation, comparative predictive performance remains informative about how effectively different predictor classes recover observed phenotypic variation under a common validation design. These studies collectively indicate that the relative predictive performance of GS and PS depends on the biological proximity of the phenomic measurement to the target trait, population structure, validation design, and trait architecture.

In our study, phenomic prediction was weaker than genomic prediction overall but its performance should be interpreted in light of the unusually long temporal and environmental separation between predictor collection and trait expression. Seedling phenotypes were collected under two distinct controlled conditions before field establishment in the fall and used to predict traits measured in subsequent growing seasons one to two years later across field sites and years. Phenomic selection based on spectral data is often most effective when predictors are collected near the time and environment of target-trait expression, where they can capture contemporaneous physiological and environmental variation associated with the trait (Krause et al., 2019; Rincent et al., 2018c). Under the temporally separated conditions evaluated here, the modest but reproducible performance of selected phenomic streams across cycles is therefore more informative than their absolute performance alone. These results do not suggest that phenomic selection can generally replace genomic selection in this breeding context. Instead, early-life stage phenotypes may support more accessible means of ranking or prioritization when precise genomic prediction is unavailable or unnecessary, particularly during early stages of domestication or population improvement before a sufficiently large genomic training population has been established. As genomic resources and accumulated training data increase, breeding programs may then transition toward greater reliance on genomic selection.

### Leaf HSV provides reproducible phenomic prediction across developmental stages and years

Among the phenomic predictors evaluated, leaf HSV consistently provided the strongest predictive signal for later-expressed field traits across both breeding cycles. The mean predictive ability of models using leaf HSV was similar across cycles, and models trained in one cycle retained comparable performance when used to predict the other. This cross-cycle transfer is important because it indicates that the predictive information captured by leaf HSV was not restricted to a single cohort, experimental year, or field site. Moreover, this transferability suggests that predictive performance was not driven solely by cycle-specific maternal-family differences or transient early-life conditioning. Although predictive performance from leaf HSV was weaker than genomic predictions on average, it approached or exceeded GS for several yield-related and reproductive traits, suggesting that early seedling color can retain information about later performance even across substantial developmental and environmental separation.

Leaf HSV may be effective because it integrates multiple aspects of plant state rather than measuring a single physiological process. Leaf color reflects variation in pigment composition, nutrient status, stress response, and developmental condition, all of which can influence growth trajectories and resource allocation (Agarwal et al., 2025; Fukano et al., 2023; Manetas, 2006; Oliveira & Santana, 2020; Sun et al., 2018; H. Zhang et al., 2022). HSV decomposition separates hue, saturation, and brightness, allowing broad chromatic differences to be represented partly independently of overall illumination and potentially aligning more closely with biologically meaningful pigment variation than raw RGB values (Yang et al., 2015). These measurements were collected during early seedling establishment, a period characterized by rapid ontogenetic changes in pigment composition, photosynthetic capacity, biomass accumulation, and resource-allocation traits (Havrilla et al., 2021). Genotypic and family-level differences in seedling growth and resource-use strategies can be pronounced during this period, making early plant condition a plausible indicator of later performance (Chybicki et al., 2025; Rowe & Leger, 2011; Rweyongeza et al., 2004; Umaña et al., 2025).

The exact biological basis of this signal remains unresolved. Leaf HSV likely reflects a mixture of genetic, maternal, developmental, and physiological variation, but the present analyses cannot determine the relative contribution of these sources. Cross-cycle transfer makes a purely cycle-specific technical or environmental explanation less likely, but it does not establish that the relevant variation is heritable, causal (Feldmann et al., 2026), or stable across more distantly related breeding populations. The strong performance of leaf HSV should therefore be interpreted as evidence that early seedling color provides a useful integrative representation of plant state, rather than as evidence for any specific molecular mechanism.

Leaf HSV strongly outperformed seed HSV, which showed little predictive value for later agronomic traits despite strong maternal-family structure, indicating that substantial biological variation in seed color did not translate into useful information for these downstream targets. This contrasts with studies in which seed-based phenomic predictors were applied to more developmentally proximal traits, including germination and establishment traits (Keaggy et al., 2025) and traits predicted from NIR spectra collected directly on intact seed material (Graciano et al., 2025). In our study, the long developmental separation between seed appearance and mature field performance likely limited the expression of useful predictive signal. Given the consistently near-zero performance across cycles and traits, seed HSV appears to have limited utility for predicting later agronomic performance in this breeding context.

Interestingly, leaf HSV outperformed the hyperspectral data streams evaluated here, despite the much greater spectral resolution of those instruments. Hyperspectral reflectance can capture fine-scale biochemical and structural variation and has been used successfully to estimate pigments, nutrients, and physiological traits in many crop systems (Grzybowski et al., 2021; Kothari et al., 2023; Paulus & Mahlein, 2020; Sarić et al., 2022; Silva-Perez et al., 2018; Yendrek et al., 2017). In this study, however, hyperspectral measurements contained substantial device-and collection-associated variation and provided weaker, more sensor-dependent prediction of mature field traits. The additional spectral detail may therefore have added information that was technically unstable, internally redundant, or poorly aligned with the downstream traits of interest (L. Xu et al., 2025). By contrast, leaf HSV may capture broad, correlated gradients in pigment status, nutrition, senescence, and stress that remain informative across development, whereas hyperspectral reflectance additionally resolves finer-scale physiological and biochemical states whose relationships with later phenotypes may be more developmentally or environmentally contingent (Magney et al., 2026; Villa et al., 2021; R. Xu et al., 2026). These results should not be interpreted as evidence that HSV is intrinsically superior to hyperspectral sensing, particularly because different hyperspectral instruments were used in the two cycles and we were unable to explore cross-cycle predictions using those sensors. Rather, across the sensors and protocols evaluated here, a simpler and more integrative color representation provided more reproducible prediction of temporally distant field traits.

### Distinct relationship structures do not guarantee predictive complementarity

Phenomic and genomic relationship matrices differed in the patterns of similarity they captured among individuals, suggesting that combining data streams might contribute nonredundant information when modeled together (Morota & Gianola, 2014). However, multi-relationship-matrix models rarely improved predictive performance beyond the stronger constituent single-relationship-matrix model. This pattern was consistent across combinations of genomic and phenomic matrices and across phenomic-only models. In particular, models combining GS and leaf HSV sometimes exceeded GS alone, but generally did not outperform the better of GS or leaf HSV considered separately. Thus, differences in relationship-matrix structure were not sufficient to produce gains in prediction. Several factors could explain this result. A second relationship matrix may represent a pattern of similarity among individuals that differs from the first, yet contribute little additional information for predicting the target trait. A second relationship matrix could also capture variation that is already represented by the stronger matrix or contribute a covariance component that is difficult to estimate precisely in the presence of other relationship structures. Accordingly, matrix dissimilarity should not be interpreted as evidence that two data streams will be predictively complementary.

These findings also reinforce the importance of choosing appropriate benchmarks for phenomic prediction models. Consistent with broader concerns about treating GS as the default comparator for PS ((F. Wang et al., 2025), multi-relationship-matrix models should be evaluated against the strongest constituent model rather than only against a single conventional baseline such as GS. These findings also suggest that the value of combining phenomic data streams should be established empirically for each prediction problem. Previous studies have shown that expanding the information represented within a phenomic relationship matrix or combining genomic and phenomic information can improve prediction under some conditions (Graciano et al., 2025; Keaggy et al., 2025; Krause et al., 2019; Rincent et al., 2025). In the present study, however, additional relationship matrices increased model complexity without providing concomitant predictive gains.

### Practical deployment of early-life stage phenomics in perennial breeding

Our results identify a realistic but bounded role for early-life stage phenomics in intermediate wheatgrass breeding. Genomic selection remained the strongest overall predictor, but leaf HSV provided modest, reproducible predictive ability across breeding cycles and transferred with little loss when models were applied to a new cycle. Because leaf HSV can be collected from inexpensive RGB images at an early developmental stage, it may be especially useful for germplasm thinning, early-stage prioritization, or identifying individuals and families that warrant more expensive genotyping or field evaluation.

The practical value of leaf HSV does not depend on replacing genomic selection. In large breeding populations, even moderate predictive ability may be useful when applied at an early decision point, particularly when the alternative is advancing all material or making selections with little information. An accessible phenomic screen could therefore reduce the number of individuals carried forward, concentrate genotyping resources on higher-priority material, or provide a preliminary ranking when genomic data are not yet available. Whether such a strategy produces net breeding value will depend on the cost of phenotyping and genotyping, the intensity and stage of selection, and the consequences of incorrectly discarding promising material.

The results also caution against equating technological sophistication with breeding utility. Across the sensors and protocols evaluated here, higher-dimensional multispectral and hyperspectral data did not consistently outperform leaf HSV, and combining relationship matrices rarely improved prediction beyond the stronger constituent model. Phenomic tools should therefore be evaluated according to the reliability, cost, portability, and decision value of the information they provide, rather than by dimensionality alone. For emerging perennial crops, targeted phenomic measurements may be most useful as an operational complement to genomic selection, not necessarily as an additional covariance term in the same model, but as a strategically deployed source of information at stages where genotyping or mature phenotyping is impractical.

### Limitations and future directions

Several limitations qualify the interpretation and deployment of these results. First, phenomic measurements were collected in a highly controlled automated growth-house environment. Although this design reduced technical and nuisance variation during data collection, it may overstate the portability of leaf HSV to breeding programs that rely on greenhouses, field nurseries, or lower-cost imaging systems. Future work should therefore further test whether leaf HSV retains predictive ability under less controlled conditions and with simpler image-acquisition protocols (Woeltjen et al., 2026). Sampling at later developmental stages may also improve prediction by reducing temporal distance from the target traits (Adak et al., 2023, 2024), but such gains would need to be weighed against the loss of early-selection value.

Second, although the imposed water dry-down treatment altered subsets of phenomic features, these changes did not translate into a consistent or functionally meaningful change in PS performance. Thus, under the controlled conditions evaluated here, deliberately increasing early-life stress did not provide an obvious advantage for phenomic prediction of later field traits. More severe, developmentally targeted, or field-representative stress treatments could still expose predictive variation not captured by the dry-down imposed here, as drought phenotyping responses are known to depend strongly on stress timing, severity, and environmental context (Tuberosa, 2012).

Third, the combined-cycle analyses require cautious interpretation. Genomic prediction improved when cycles were pooled. Published IWG genomic predictions based on several thousand genets have generally exceeded the within-cycle model accuracies observed here, suggesting that models trained on approximately 1,000 individuals per cycle may not yet have reached their performance ceiling (Bajgain et al., 2025; Crain, DeHaan, et al., 2021b). In contrast, pooling cycles did not consistently improve phenomic prediction: CropReporter performance remained largely unchanged, whereas the large apparent increase in leaf HSV performance at least partly reflected residual cycle-associated structure that remained despite adjustment. This divergence does not establish that phenomic models are saturated at the present sample size; substantially larger, consistently measured training populations may be needed to determine whether weaker phenomic signals can be estimated more reliably. Different hyperspectral instruments were also used in the two cycles, limiting direct sensor comparisons and preventing straightforward pooling of those data streams. Replication using consistent instrumentation, explicit calibration-transfer methods, and independent breeding populations will be necessary to determine how broadly these results generalize.

Finally, we used a standardized relationship-matrix modeling framework to compare data streams rather than exhaustively tuning algorithms, kernel architecture, or preprocessing strategies (Montesinos-López et al., 2021b). The reported differences therefore reflect performance under a common analytical framework rather than the maximum achievable accuracy of each data stream. Beyond this analytical limitation, the biological basis of the strongest phenomic signal also remains unresolved. Future work should therefore evaluate both alternative modeling strategies and the extent to which leaf HSV prediction reflects heritable variation, persistent developmental state, maternal effects, or other plant-level processes. Models that explicitly retain environmental and temporal interactions may further distinguish predictors of stable performance from those that capture context-specific responses.

## Data availability

The sequencing data generated for this study are available through the NCBI Sequence Read Archive under BioProject accession PRJNA1044453. Phenotypic data and the code of record used to generate the analyses and figures are available through Figshare at doi.org/10.6084/m9.figshare.33354318. Analysis scripts are also available through GitHub at https://github.com/znh1992/kernza_ps.

## Author contributions

**Zachary N. Harris**: Data curation; formal analysis; methodology; visualization; writing—original draft; writing—review and editing. **Jackson Braley**: Data curation; investigation; methodology. **Eric Cassetta**: Data curation; investigation; methodology. **Jared L. Crain**: Data curation; formal analysis; investigation; methodology; writing—review and editing. **Lee R. DeHaan**: Conceptualization; data curation; formal analysis; funding acquisition; investigation; methodology; project administration; resources; supervision; writing—review and editing. **David L. Van Tassel**: Conceptualization; funding acquisition; project administration; supervision; writing—review and editing. **Allison J. Miller**: Conceptualization; funding acquisition; investigation; project administration; resources; supervision; writing—review and editing. **Matthew J. Rubin**: Conceptualization; data curation; formal analysis; funding acquisition; investigation; methodology; project administration; supervision; visualization; writing—original draft; writing—review and editing.

## Conflict of Interest

The authors declare no conflicts of interest.

## Declaration of Generative AI and AI-Assisted Technologies

During preparation of this work, the authors used OpenAI ChatGPT (primarily GPT-5.6 Sol) for language editing and manuscript organization. The authors reviewed and edited all AI-assisted content and take full responsibility for the published article.

## Acknowledgements

We thank the Foundation for Food and Agriculture (FFAR; CA20-SS-0000000123 and 24-001143), The Perennial Agriculture Project in conjunction with the Malone Family Land Preservation Foundation and The Land Institute, Donald Danforth Plant Science Center, The Land Institute, Taylor Geospatial Institute, and Saint Louis University for funding this work. We express our gratitude to John and Pauline Cella of Planthaven Farms for access to land for this study. We especially thank Brandon Schlautman, Jesse Poland, Kathryn Turner, and Luis Diaz-Garcia for contributions to project conceptualization and funding acquisition; Jenna Hershberger for assistance with data collection and manuscript review; Tyler Thrash for data collection and curation; and Noah Fahlgren, Jorge Gutierrez, and Haley Schuhl for assistance with data collection and methodology. We also thank members of the Miller Lab, Rubin Lab, The Danforth Plant Science Center, and The Land Institute for their extensive assistance with planting, plant care, field and controlled-environment phenotyping, and data collection. Finally, we thank the Danforth Center Plant Growth Facilities (RRID:SCR_024902), Danforth Center Field Research Site (RRID:SCR_027897) Danforth Center Phenotyping Facility (RRID:SCR_019049), and Danforth Center Data Science Facility (RRID:SCR_027573) for their support of this work.

## Abbreviations

AR1 × AR1: first-order autoregressive spatial correlation in two dimensions
BLUP: best linear unbiased prediction
DPI: dots per inch
F0: minimum chlorophyll fluorescence
Fv/Fm: maximum quantum efficiency of photosystem II
GRM: genomic relationship matrix
GS: genomic selection
HSV: hue, saturation, value
IWG: intermediate wheatgrass
KS: Kansas field environment
NIR: near-infrared
NDVI: normalized difference vegetation index
PCA: principal component analysis
PS: phenomic selection
PVE: proportion of variance explained
QTL: quantitative trait locus
RGB: red, green, blue
RKHS: reproducing kernel Hilbert space
SNP: single nucleotide polymorphism
STL: St. Louis field environment
VWC: volumetric water content

## Supplemental Information

**Supplemental Table 1.** Field-trait descriptions and corresponding standardized trait database identifiers used in this study.

**Supplemental Table 2.** Brown-Forsythe summary tables for each of the predictor data streams across water treatments.

**Supplemental Table 3.** Analysis of variance tables for field traits, including model terms, degrees of freedom, sums of squares, test statistics, p-values, and estimated proportion of variance explained (PVE). These data underlie Figure 2.

**Supplemental Table 4.** Genomic selection and phenomic selection performance for each trait, relationship matrix, breeding cycle, and cross-validation fold across the six data streams. Mean and standard deviation of predictive performance were calculated for each trait-by-relationship-matrix-by-cycle combination. These data underlie Figure 3.

**Supplemental Table 5.** Predictive performance of multi-relationship-matrix models for each trait, breeding cycle, and cross-validation fold, with corresponding mean and standard deviation summaries. These data partially underlie Figure 4.

**Supplemental Table 6.** Genomic and phenomic prediction performance within and across breeding cycles, including models evaluated within Cycle 11, within Cycle 12, from Cycle 11 into Cycle 12, from Cycle 12 into Cycle 11, and in the combined Cycle 11+Cycle 12 dataset. Mean performance is reported for analyses using k-fold cross-validation; cross-cycle predictions consist of a single train-test evaluation.

**Supplemental Figure 1:**
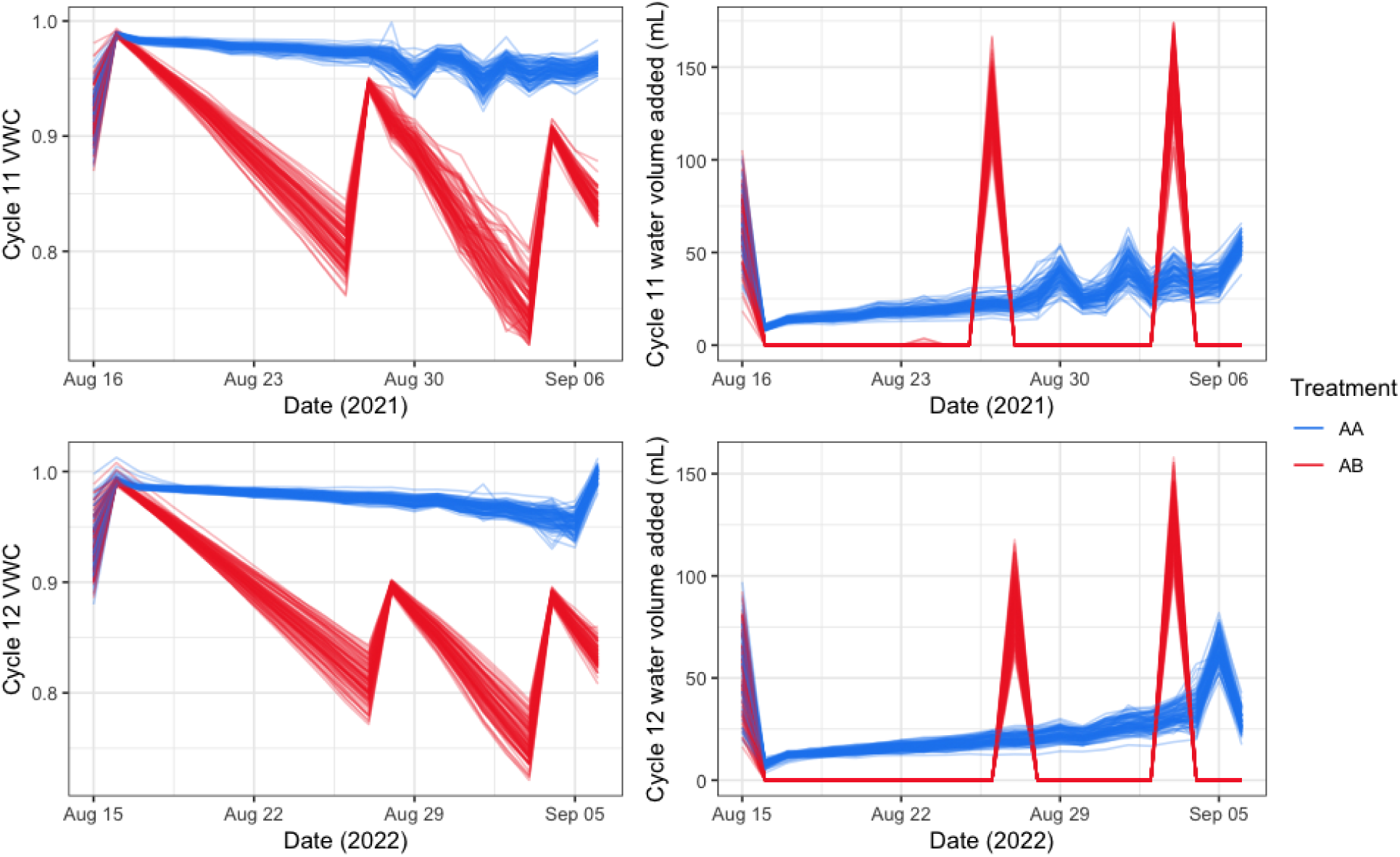
Water treatment regimes for LemnaTec growth. Watering treatments were applied such that well-watered plants (treatment AA) were watered to nearly constant daily mass. Water dry-down plants (Treatment AB) were allowed to dry down to 70-80% volumetric water content (VWC) before being watered back to initial weight (left panels). Right panels show how much water was added per day. For simplicity, lines show averaged families across cycles.

**Supplemental Figure 2.**
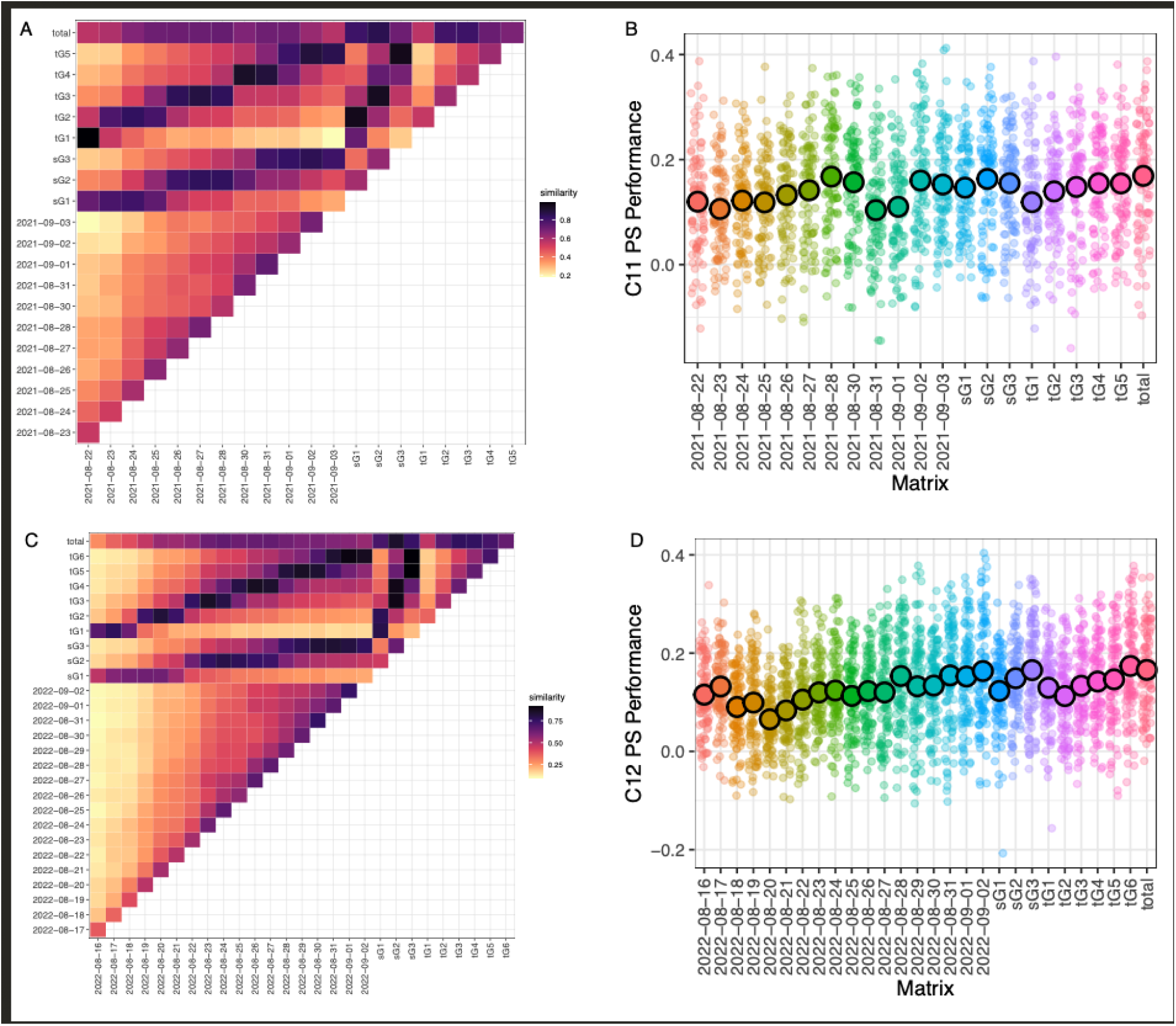
Decision context for selecting the leaf HSV imaging window in Cycle 11 and Cycle 12. (A, C) Mantel-like comparisons of leaf HSV relationship matrices constructed from individual imaging days, grouped imaging windows, and all imaging days combined for Cycle 11 (A) and Cycle 12 (C). Daily matrices are labeled by collection date; six-day and three-day groupings are denoted by sG and tG, respectively, and “total” represents the relationship matrix constructed using all available imaging days. Cell color indicates the correlation between pairs of relationship matrices; no permutation or bootstrap procedure was used to assess statistical significance. (B, D) Within-cycle phenomic selection performance for the corresponding leaf HSV relationship matrices in Cycle 11 (B) and Cycle 12 (D). Each translucent point represents the Pearson correlation between observed and predicted values for one trait in one cross-validation fold. Filled circles show mean predictive performance across traits and folds, with error bars denoting 83% confidence intervals. Predictive performance varied modestly among individual dates and grouped imaging windows, supporting the use of measurements collected near the end of the imaging period in subsequent analyses.

**Supplemental Figure 3.**
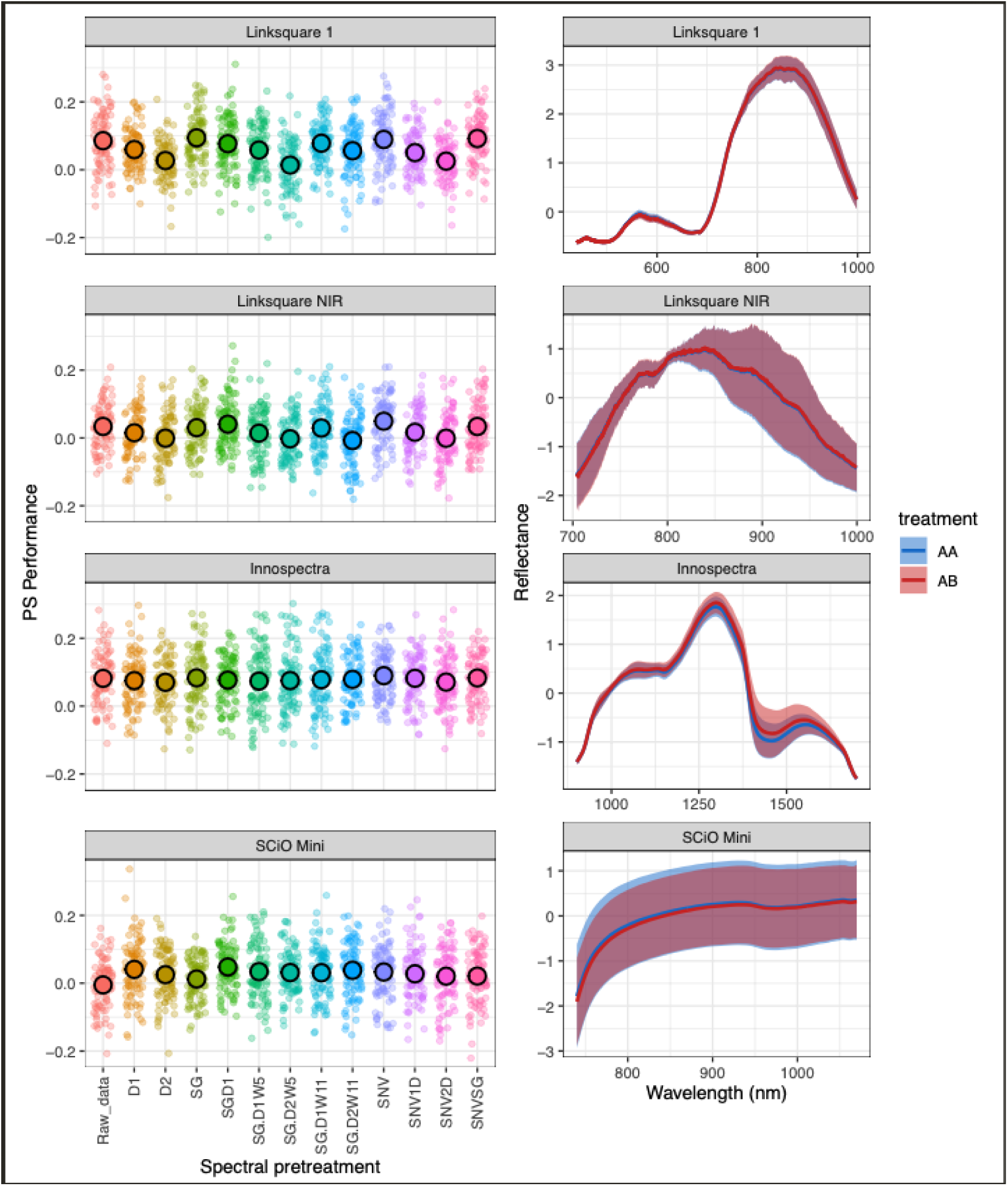
Decision context for hyperspectral preprocessing in Cycle 11 and Cycle 12. For each sensor, the left panel shows within-cycle phenomic selection performance across the evaluated spectral pretreatments, and the right panel shows raw hyperspectral reflectance profiles for the two seedling watering treatments. Each translucent point in the performance panels represents the Pearson correlation between observed and predicted values for one trait in one cross-validation fold. Filled circles show mean predictive performance across traits and folds, with error bars denoting 83% confidence intervals. Reflectance differences between watering treatments were generally modest relative to the overall spectral structure, and predictive performance varied little among pretreatments, supporting the use of untransformed reflectance in subsequent analyses.

**Supplemental Figure 4.**
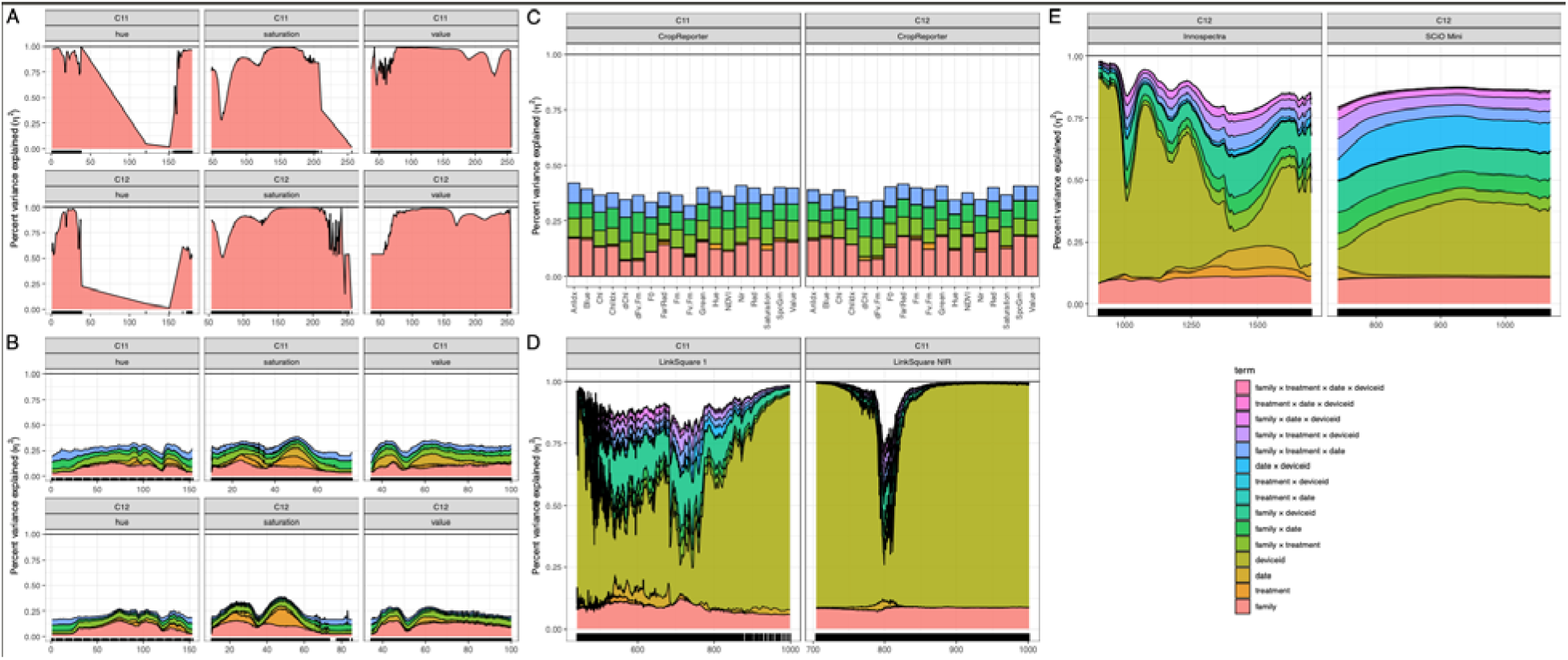
Decomposition of variance explained by maternal family, watering treatment, and stream-specific technical effects across phenomic predictors in breeding cycles 11 and 12. (A) Seed HSV, (B) Leaf HSV, (C) CropReporter, (D) Cycle 11 hyperspectral sensors, and (E) Cycle 12 hyperspectral sensors. Colored regions show the proportion of variance explained by each modeled effect for individual predictor features. Technical effects and interactions varied among phenomic streams according to their respective sampling and measurement designs. Where present, black tick marks along the x-axis indicate individual predictor features. Seed HSV was measured before assignment to watering treatment and was therefore modeled only for maternal-family effects.

**Supplemental Figure 5.**
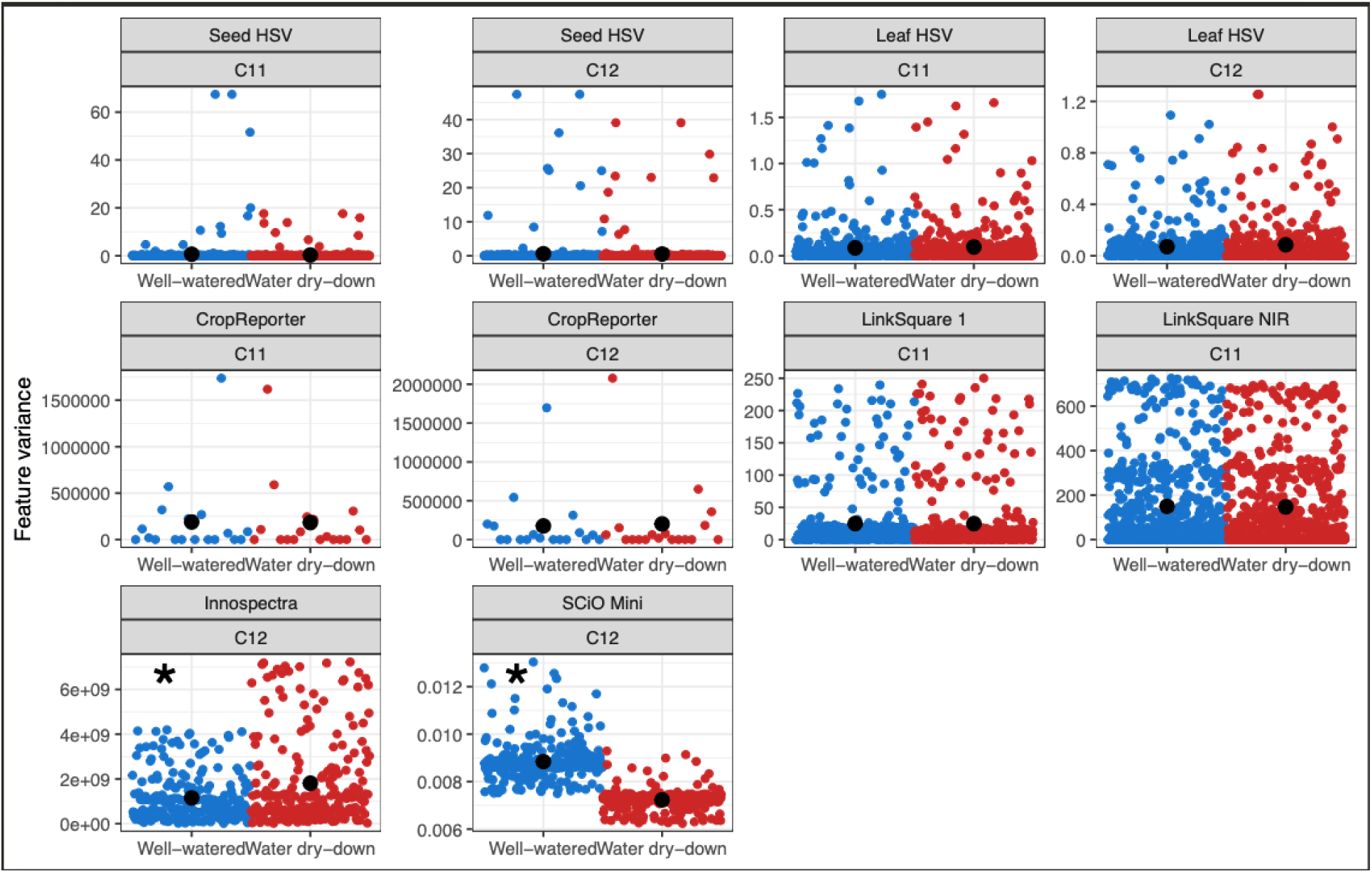
Distribution of feature-level variance across watering treatments for each phenomic data stream and breeding cycle. Points represent the variance of individual predictor features, calculated separately for well-watered and water dry-down plants and expressed in feature-specific units. Black filled circles indicate the mean feature variance within each treatment. Asterisks denote data streams for which mean feature variance differed between watering treatments. Across most data streams and cycles, variance distributions were similar between treatments; significant differences were detected only for the Cycle 12 hyperspectral sensors.

**Supplemental Figure 6.**
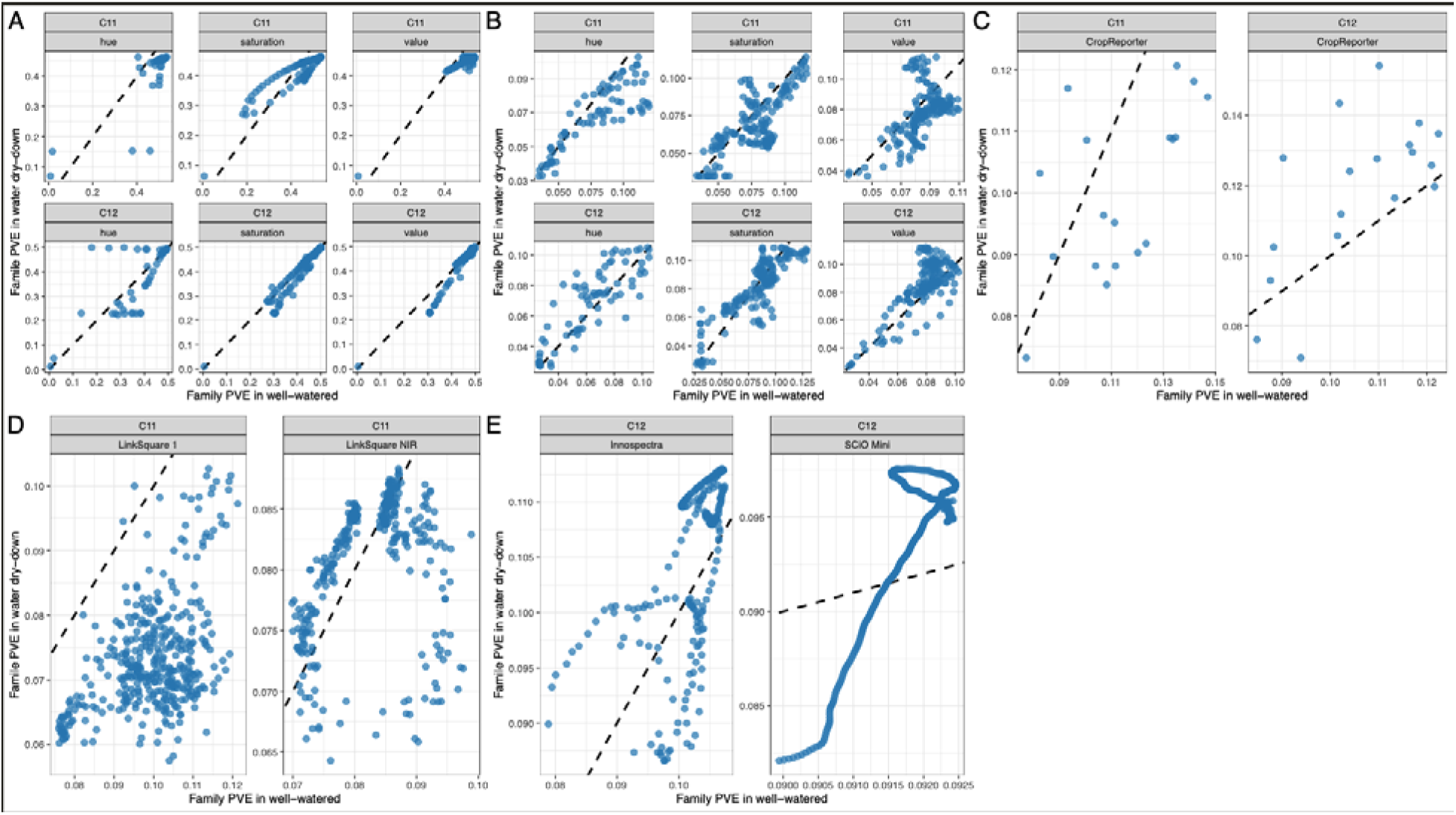
Maternal-family variance explained within well-watered and water dry-down treatments across phenomic data streams and breeding cycles. Points represent individual predictor features, with maternal-family PVE estimated separately within each watering treatment. The dashed line indicates a 1:1 relationship; points above the line have greater maternal-family PVE under water dry-down, whereas points below the line have greater maternal-family PVE under well-watered conditions. Panels show (A) seed HSV, (B) leaf HSV, (C) CropReporter, (D) Cycle 11 hyperspectral sensors, and (E) Cycle 12 hyperspectral sensors. Across data streams, watering treatment did not produce a consistent directional shift in the proportion of phenomic variance attributable to maternal family.

**Supplemental Figure 7.**
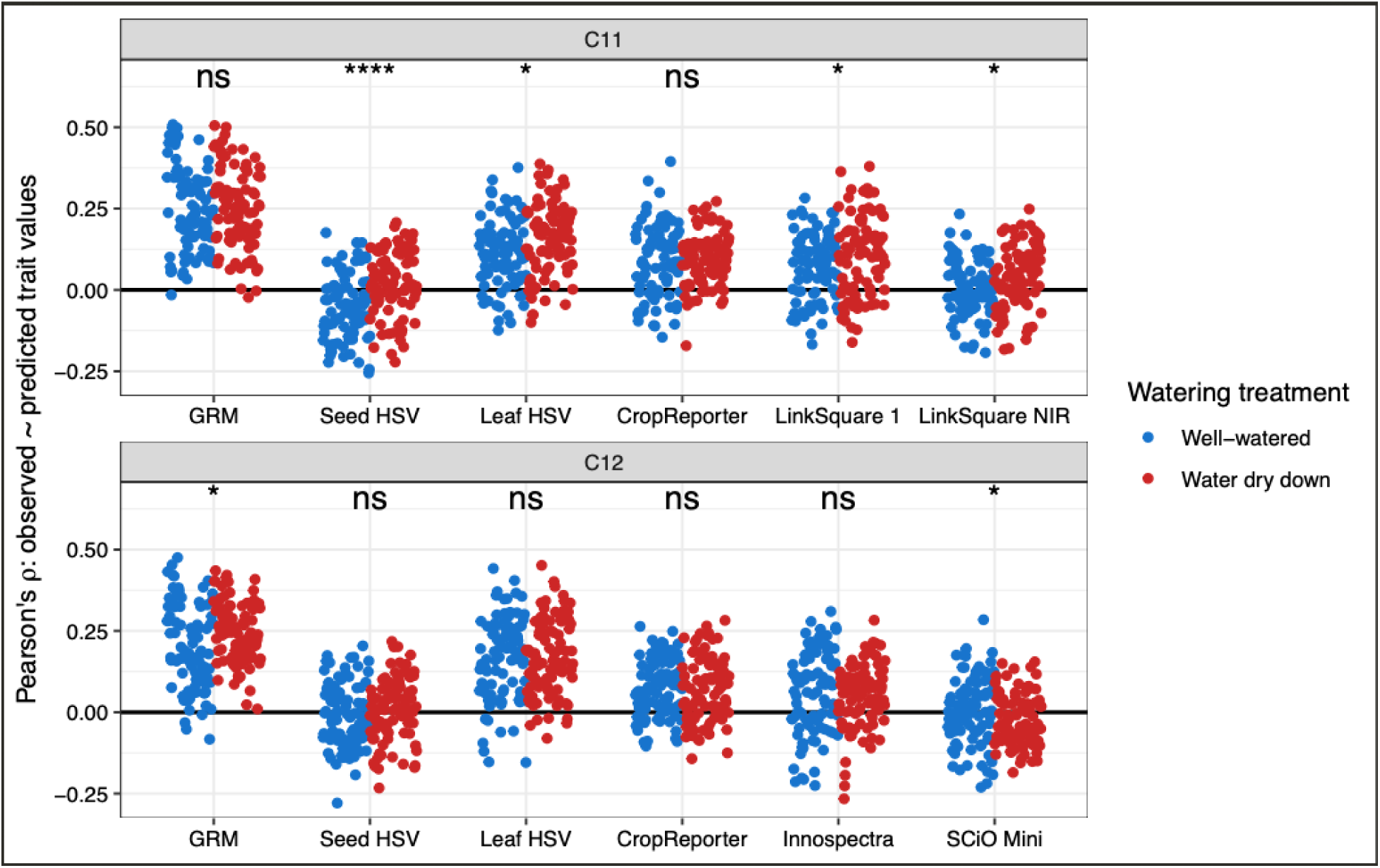
Phenomic and genomic prediction performance within watering treatments across breeding cycles. Points represent Pearson correlations between observed and predicted field-trait values for individual trait-by-fold combinations, estimated separately for well-watered and water dry-down plants. Predictive performance is shown for the genomic relationship matrix (GRM) and relationship matrices derived from each phenomic data stream available within Cycle 11 and Cycle 12. Significance annotations indicate tests for differences in predictive performance between watering treatments within each predictor stream (*p* < 0.05, *; *p* < 0.01, **; *p* < 0.005, ***; *p* < 0.001, ****). Although several comparisons were statistically significant, the corresponding effect sizes were generally small and did not alter the functional interpretation of model performance, particularly where correlations remained near zero in both treatments.

**Supplemental Figure 8:**
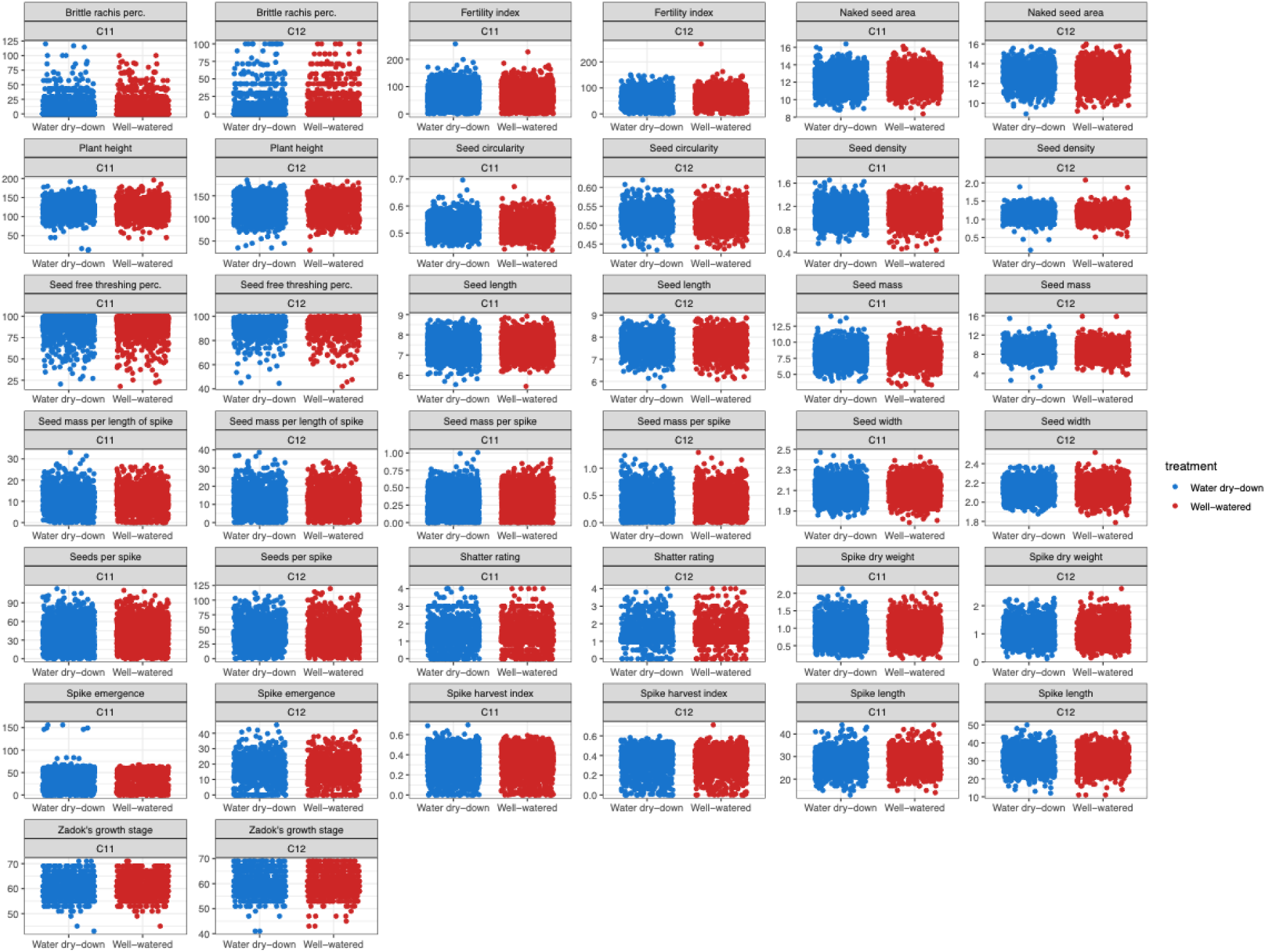
Field-trait distributions by seedling watering treatment and breeding cycle. Points represent individual field phenotypes for traits measured in Cycle 11 and Cycle 12, grouped according to the well-watered or water dry-down treatment imposed during seedling growth. Across traits, distributions were broadly similar between watering treatments. Formal trait-by-trait tests detected no watering-treatment effects that remained significant after Benjamini–Hochberg correction for multiple testing.

**Supplemental Figure 9.**
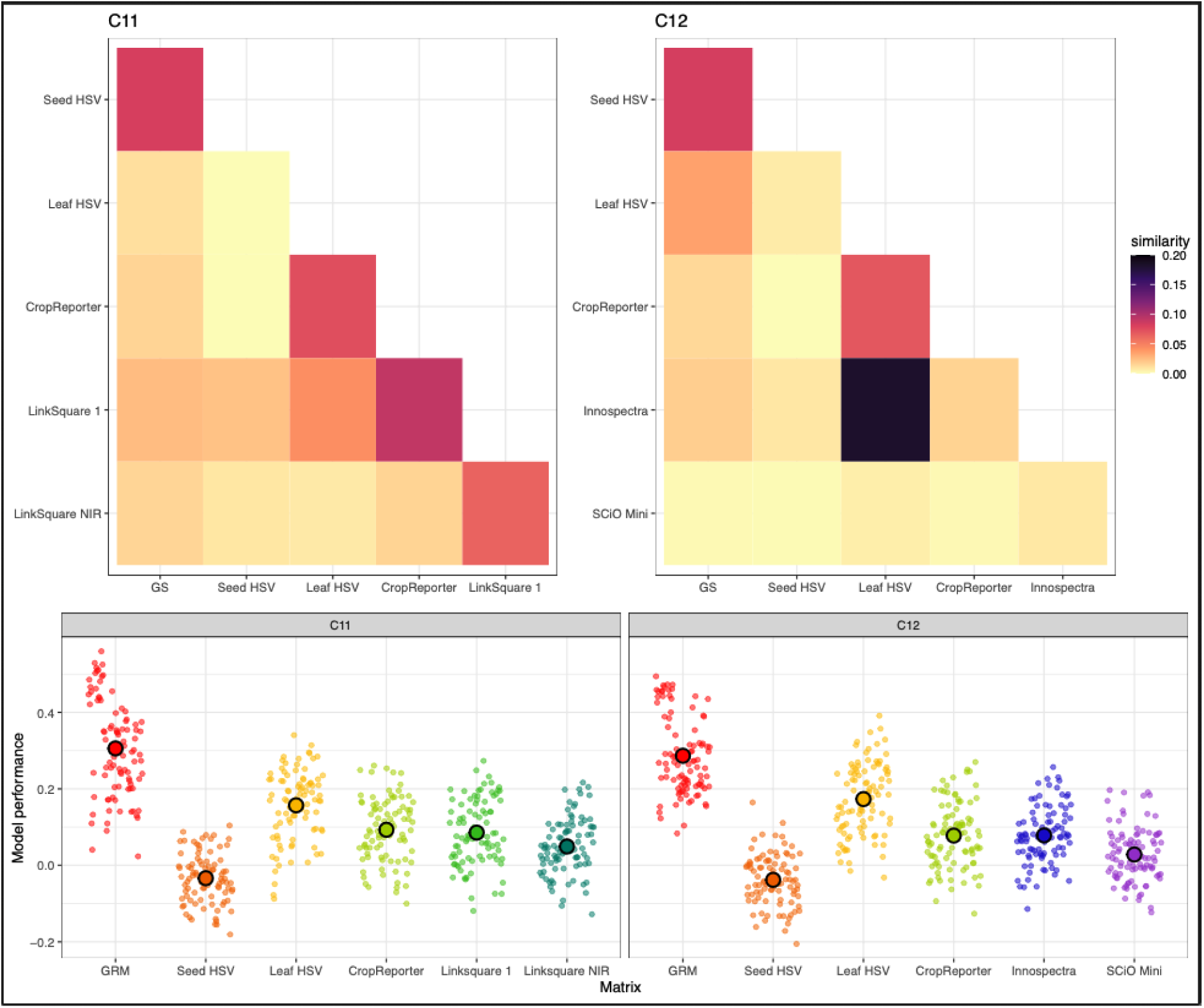
Similarity and predictive performance of genomic and phenomic relationship matrices across Cycle 11 and Cycle 12. (Top panels) Mantel-like similarity comparisons of the genomic relationship matrix (GRM) and relationship matrices derived from each phenomic data stream in Cycle 11 and Cycle 12. Cell color indicates the correlation between the individual-by-individual relationship structures represented by each pair of matrices; no permutation or bootstrap procedure was used to assess statistical significance. (Bottom panels) Within-cycle predictive performance of single-relationship-matrix models in Cycle 11 and Cycle 12. Predictors included the GRM, seed HSV, leaf HSV, CropReporter traits, and cycle-specific hyperspectral data from LinkSquare 1 and LinkSquare NIR in Cycle 11 and Innospectra and SCiO Mini in Cycle 12. Each small translucent point represents the Pearson correlation between observed and predicted values for one trait in one cross-validation fold. Larger, black-outlined circles show mean predictive performance across traits and folds, with error bars (often hidden behind the points) denoting 83% confidence intervals. Across both cycles, the GRM provided the strongest overall prediction, leaf HSV was the strongest phenomic predictor, and seed HSV showed little predictive ability despite differences in relationship structure among data streams.

**Supplemental Figure 10.**
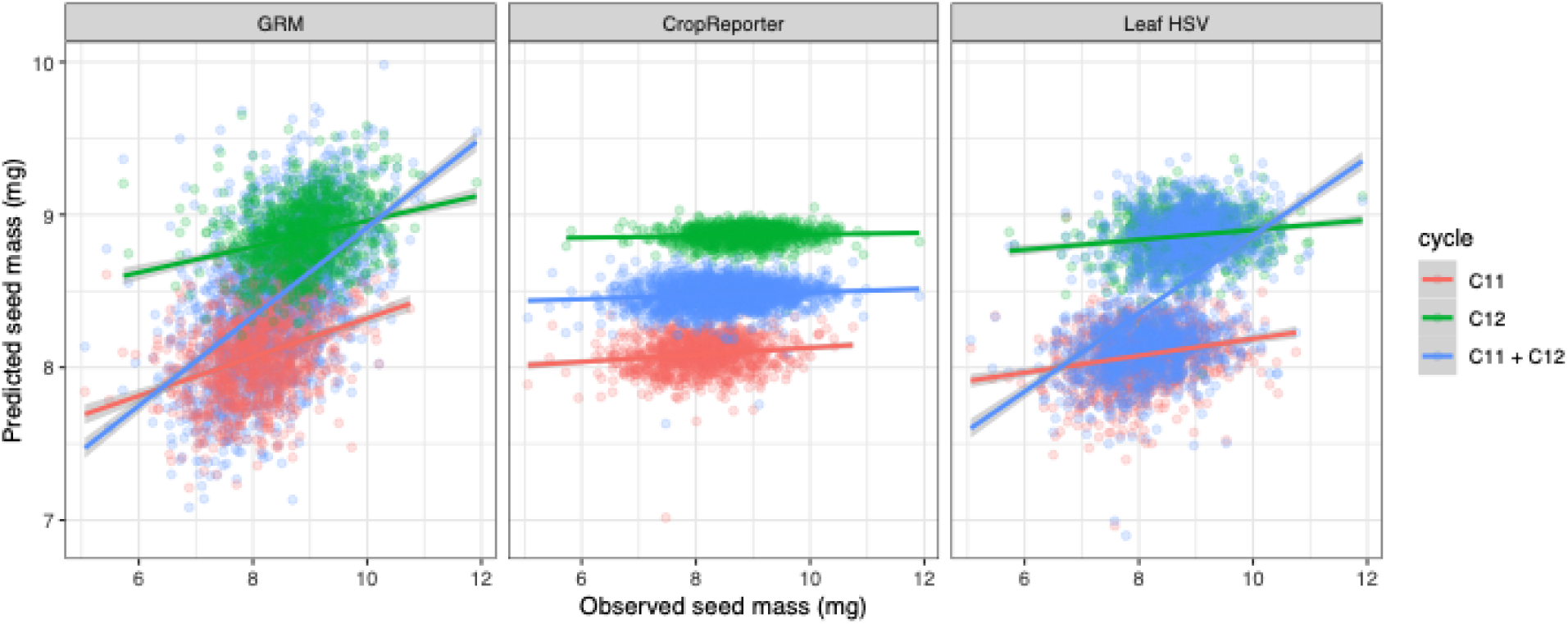
Residual cycle structure inflates the apparent predictive performance of combined-cycle Leaf HSV models. Observed and predicted seed mass are shown for models based on the genomic relationship matrix (GRM), CropReporter phenotypes, and Leaf HSV phenotypes. Points are colored by breeding cycle, and lines show linear relationships fitted separately for Cycle 11, Cycle 12, and the combined Cycle 11+Cycle 12 dataset; shaded regions indicate confidence intervals. GRM predictions retain clear within-cycle relationships, and their improved performance in the combined-cycle analysis is therefore consistent with increased training information rather than an aggregation artifact. For CropReporter, the across-cycle adjustment procedure successfully removed most cycle-level displacement, and the pooled relationship remains consistent with the weak relationships observed within each cycle. In contrast, Leaf HSV predictions retain substantial separation between Cycle 11 and Cycle 12 despite the same attempt to control cycle effects. Consequently, the steep combined-cycle relationship is driven largely by between-cycle differences in observed and predicted seed mass rather than by improved within-cycle prediction. The apparent increase in Leaf HSV predictive ability after combining cycles cannot be attributed cleanly to increased training population size and might be an artifact of unresolved cycle structure.

## Notes

### Competing Interest Statement

The authors have declared no competing interest.

## References

Adak, A., DeSalvio, A. J., Arik, M. A., & Murray, S. C. (2024). Field-based high-throughput phenotyping enhances phenomic and genomic predictions for grain yield and plant height across years in maize. G3 (Bethesda, Md.), 14(7). 10.1093/g3journal/jkae092

Adak, A., Murray, S. C., & Anderson, S. L. (2023). Temporal phenomic predictions from unoccupied aerial systems can outperform genomic predictions. G3 (Bethesda, Md.), 13(1). 10.1093/g3journal/jkac294

Adunola, P., Ferrão, M. A. G., Ferrão, R. G., da Fonseca, A. F. A., Volpi, P. S., Comério, M., Verdin Filho, A. C., Munoz, P. R., & Ferrão, L. F. V. (2023). Genomic selection for genotype performance and environmental stability in Coffea canephora. G3 (Bethesda, Md.), 13(6). 10.1093/g3journal/jkad062

Adunola, P., Tavares Flores, E., Azevedo, C., Casorzo, G., Ghimire, L., Ferrão, L. F. V., & Munoz, P. R. (2024). Phenomic assisted selection: Assessment of the potential of near infrared spectroscopy for blueberry breeding. Plant Phenome Journal, 7(1). 10.1002/ppj2.70010

Agarwal, A., de Jesus Colwell, F., Correa Galvis, V. A., Hill, T. R., Boonham, N., & Prashar, A. (2025). Assessing nutritional pigment content of green and red leafy vegetables by image analysis: Catching the “red herring” of plant digital color processing via machine learning. Biology Methods & Protocols, 10(1), bpaf027.

Altendorf, K. R., DeHaan, L. R., & Anderson, J. (2022). Genetic architecture of yield component traits in the new perennial grain crop, intermediate wheatgrass. Crop Science, 62(2), 880–892.

Asbjornsen, H., Hernandez-Santana, V., Liebman, M., Bayala, J., Chen, J., Helmers, M., Ong, C. K., & Schulte, L. A. (2014). Targeting perennial vegetation in agricultural landscapes for enhancing ecosystem services. Renewable Agriculture and Food Systems, 29(2), 101–125.

Bajgain, P., Crain, J. L., Cattani, D. J., Larson, S. R., Altendorf, K. R., Anderson, J. A., Crews, T. E., Hu, Y., Poland, J. A., Turner, M. K., Westerbergh, A., & DeHaan, L. R. (2022). Breeding Intermediate Wheatgrass for Grain Production. In Plant Breeding Reviews (pp. 119–217). Wiley. 10.1002/9781119874157.ch3

Bajgain, P., Jungers, J. M., & Anderson, J. A. (2024). Genetic constitution and variability in synthetic populations of intermediate wheatgrass, an outcrossing perennial grain crop. G3 (Bethesda, Md.), 14(9). 10.1093/g3journal/jkae154

Bajgain, P., Stoll, H., & Anderson, J. A. (2025). Improving complex agronomic and domestication traits in the perennial grain crop intermediate wheatgrass with genetic mapping and genomic prediction. The Plant Genome, 18(1), e20498.

Brault, C., Segura, V., This, P., Le Cunff, L., Flutre, T., François, P., Pons, T., Péros, J.-P., & Doligez, A. (2022). Across-population genomic prediction in grapevine opens up promising prospects for breeding. Horticulture Research, 9, uhac041.

Browning, B. L., & Browning, S. R. (2016). Genotype Imputation with Millions of Reference Samples. American Journal of Human Genetics, 98(1), 116–126.

Chandel, N. S., Tiwari, P. S., Singh, K. P., Jat, D., Gaikwad, B. B., Tripathi, H., & Golhani, K. (2019). Yield Prediction in Wheat (*Triticum aestivum* L.) using Spectral Reflectance Indices. Current Science, 116(2), 272.

Chapman, E. A., Thomsen, H. C., Tulloch, S., Correia, P. M. P., Luo, G., Najafi, J., DeHaan, L. R., Crews, T. E., Olsson, L., Lundquist, P.-O., Westerbergh, A., Pedas, P. R., Knudsen, S., & Palmgren, M. (2022). Perennials as Future Grain Crops: Opportunities and Challenges. Frontiers in Plant Science, 13, 898769.

Chybicki, I. J., Suszka, J., Meyza, K., & Iszkuło, G. (2025). Heritability of early seedling growth in Taxus baccata, a declining conifer. Scientific Reports, 15(1), 35214.

Covarrubias-Pazaran, G. (2016). Genome-Assisted Prediction of Quantitative Traits Using the R Package sommer. PloS One, 11(6), e0156744.

Crain, J., Bajgain, P., Anderson, J., Zhang, X., DeHaan, L., & Poland, J. (2020). Enhancing Crop Domestication Through Genomic Selection, a Case Study of Intermediate Wheatgrass. Frontiers in Plant Science, 11, 319.

Crain, J., DeHaan, L., & Poland, J. (2021a). Genomic prediction enables rapid selection of high-performing genets in an intermediate wheatgrass breeding program. The Plant Genome, 14(2), e20080.

Crain, J., DeHaan, L., & Poland, J. (2021b). Genomic prediction enables rapid selection of high-performing genets in an intermediate wheatgrass breeding program. The Plant Genome, 14(2), e20080.

Crain, J., Haghighattalab, A., DeHaan, L., & Poland, J. (2021a). Development of whole-genome prediction models to increase the rate of genetic gain in intermediate wheatgrass (Thinopyrum intermedium) breeding. The Plant Genome, 14(2), e20089.

Crain, J., Haghighattalab, A., DeHaan, L., & Poland, J. (2021b). Development of whole-genome prediction models to increase the rate of genetic gain in intermediate wheatgrass (Thinopyrum intermedium) breeding. The Plant Genome, 14(2), e20089.

Crain, J., Larson, S., Dorn, K., DeHaan, L., & Poland, J. (2022). Genetic architecture and QTL selection response for Kernza perennial grain domestication traits. TAG. Theoretical and Applied Genetics. Theoretische Und Angewandte Genetik, 135(8), 2769–2784.

Crain, J., Wagoner, P., Larson, S., & DeHaan, L. (2024). Origin of current intermediate wheatgrass germplasm being developed for Kernza grain production. Genetic Resources and Crop Evolution, 71(8), 4963–4978.

Culman, S., Pinto, P., Pugliese, J., Crews, T., DeHaan, L., Jungers, J., Larsen, J., Ryan, M., Schipanski, M., Sulc, M., Wayman, S., Wiedenhoeft, M., Stoltenberg, D., & Picasso, V. (2023). Forage harvest management impacts “Kernza” intermediate wheatgrass productivity across North America. Agronomy Journal, 115(5), 2424–2438.

DeHaan, L. R., Wang, S., Larson, S. R., Cattani, D. J., Zhang, X., & Kantarski, T. (2014). Current efforts to develop perennial wheat and domesticate Thinopyrum intermedium as a perennial grain. Perennial Crops for Food Security Proceedings of the FAO Expert Workshop, 72–89.

de Verdal, H., Segura, V., Pot, D., Salas, N., Garin, V., Rakotoson, T., Raboin, L.-M., VomBrocke, K., Dusserre, J., Castro Pacheco, S. A., & Grenier, C. (2024). Performance of phenomic selection in rice: Effects of population size and genotype-environment interactions on predictive ability. PloS One, 19(12), e0309502.

Eastburn, D. J., Roche, L. M., Doran, M. P., Blake, P. R., Bouril, C. S., Gamble, G., & Gornish, E. S. (2018). Seeding plants for long-term multiple ecosystem service goals. Journal of Environmental Management, 211, 191–197.

Edwards, S. M., Buntjer, J. B., Jackson, R., Bentley, A. R., Lage, J., Byrne, E., Burt, C., Jack, P., Berry, S., Flatman, E., Poupard, B., Smith, S., Hayes, C., Gaynor, R. C., Gorjanc, G., Howell, P., Ober, E., Mackay, I. J., & Hickey, J. M. (2019). The effects of training population design on genomic prediction accuracy in wheat. TAG. Theoretical and Applied Genetics. Theoretische Und Angewandte Genetik, 132(7), 1943–1952.

Endelman, J. B. (2011). Ridge regression and other kernels for genomic selection with R package rrBLUP. The Plant Genome, 4(3), 250–255.

Esfandyari, H., Fè, D., Tessema, B. B., Janss, L. L., & Jensen, J. (2020). Effects of Different Strategies for Exploiting Genomic Selection in Perennial Ryegrass Breeding Programs. G3 (Bethesda, Md.), 10(10), 3783–3795.

Feldmann, M. J., Wang, F., & Runcie, D. E. (2026). Philosophy of phenomic prediction and its incompatibility with causal inference. Plant Phenome Journal, 9(1). 10.1002/ppj2.70091

Fritsche-Neto, R., Resende, R. T., Olivoto, T., Garcia-Abadillo, J., Nascimento, M., Bahia, M. A. M., Jarquin, D., & Vieira, R. A. (2025). Prediction based breeding: Modern tools to optimize and reshape programs. Crop Science, 65(5). 10.1002/csc2.70175

Fukano, Y., Yamori, W., Misu, H., Sato, M. P., Shirasawa, K., Tachiki, Y., & Uchida, K. (2023). From green to red: Urban heat stress drives leaf color evolution. Science Advances. 10.1126/sciadv.abq3542

Gilmour, A. R., Gogel, B. J., Cullis, B. R., Welham, S. J., Thompson, R., Butler, D., Cherry, M., Collins, D., Dutkowski, G., Harding, S. A., & Others. (2015). ASReml User Guide Release 4.1 Structural Specification.

Glaubitz, J. C., Casstevens, T. M., Lu, F., Harriman, J., Elshire, R. J., Sun, Q., & Buckler, E. S. (2014). TASSEL-GBS: a high capacity genotyping by sequencing analysis pipeline. PloS One, 9(2), e90346.

Graciano, R. P., Peixoto, M. A., Leach, K. A., Suzuki, N., Gustin, J. L., Settles, A. M., Armstrong, P. R., & Resende, M. F. R., Jr. (2025). Integrating phenomic selection using single-kernel near-infrared spectroscopy and genomic selection for corn breeding improvement. TAG. Theoretical and Applied Genetics. Theoretische Und Angewandte Genetik, 138(3), 60.

Grzybowski, M., Wijewardane, N. K., Atefi, A., Ge, Y., & Schnable, J. C. (2021). Hyperspectral reflectance-based phenotyping for quantitative genetics in crops: Progress and challenges. Plant Communications, 2(4), 100209.

Havrilla, C. A., Munson, S. M., Yackulic, E. O., & Butterfield, B. J. (2021). Ontogenetic trait shifts: Seedlings display high trait variability during early stages of development. Functional Ecology, 35(11), 2409–2423.

Hershberger, J., Morales, N., Simoes, C. C., Ellerbrock, B., Bauchet, G., Mueller, L. A., & Gore, M. A. (2021). Making waves in Breedbase: An integrated spectral data storage and analysis pipeline for plant breeding programs. The Plant Phenome Journal, 4(1). 10.1002/ppj2.20012

Jackson, R., Buntjer, J. B., Bentley, A. R., Lage, J., Byrne, E., Burt, C., Jack, P., Berry, S., Flatman, E., Poupard, B., Smith, S., Hayes, C., Barber, T., Love, B., Gaynor, R. C., Gorjanc, G., Howell, P., Mackay, I. J., Hickey, J. M., & Ober, E. S. (2023). Phenomic and genomic prediction of yield on multiple locations in winter wheat. Frontiers in Genetics, 14, 1164935.

Jackson, W. (1980). New Roots for Agriculture. U of Nebraska Press.

Johansen, N. H., Bellucci, A., Hansen, P. B., Marum, P., Amdahl, H., Gylstrøm, K. H., Rognli, O. A., Kemešytė, V., Brazauskas, G., Greve, M., Persson, C., Isolahti, M., Helgadóttir, Á., Aavola, R., Asp, T., & Ramstein, G. P. (2025). Genomic prediction of agronomic traits in perennial ryegrass (Lolium perenne L.) and genotype x environment interactions at the limit of the species distribution. TAG. Theoretical and Applied Genetics. Theoretische Und Angewandte Genetik, 138(11), 281.

Jungers, J. M., Schiffner, S., Sheaffer, C., Ehlke, N. J., DeHaan, L., Torrion, J., Noland, R. L., & Franco, J. G. (2022). Effects of seeding date on grain and biomass yield of intermediate wheatgrass. Agronomy Journal, 114(4), 2342–2351.

Jung, M., Hodel, M., Knauf, A., Kupper, D., Neuditschko, M., Bühlmann-Schütz, S., Studer, B., Patocchi, A., & Broggini, G. A. (2025). Evaluation of genomic and phenomic prediction for application in apple breeding. BMC Plant Biology, 25(1), 103.

Keaggy, W., Harris, Z. N., Braley, J., Cassetta, E., Gutierrez, J., Piotter, E., Schuhl, H., Crain, J. L., DeHaan, L. R., Turner, M. K., Fahlgren, N., Hershberger, J., Schlautman, B., Van Tassel, D. L., Rubin, M. J., & Miller, A. J. (2025). Leveraging high dimensional seed colorspace in phenomic selection models for herbaceous perennial crops. Plant Phenome Journal, 8(1). 10.1002/ppj2.70045

Kothari, S., Beauchamp-Rioux, R., Laliberté, E., & Cavender-Bares, J. (2023). Reflectance spectroscopy allows rapid, accurate and non destructive estimates of functional traits from pressed leaves. Methods in Ecology and Evolution, 14(2), 385–401.

Krause, M. R., González-Pérez, L., Crossa, J., Pérez-Rodríguez, P., Montesinos-López, O., Singh, R. P., Dreisigacker, S., Poland, J., Rutkoski, J., Sorrells, M., Gore, M. A., & Mondal, S. (2019). Hyperspectral Reflectance-Derived Relationship Matrices for Genomic Prediction of Grain Yield in Wheat. G3 (Bethesda, Md.), 9(4), 1231–1247.

Lane, H. M., Murray, S. C., Montesinos López, O. A., Montesinos López, A., Crossa, J., Rooney, D. K., Barrero-Farfan, I. D., De La Fuente, G. N., & Morgan, C. L. S. (2020). Phenomic selection and prediction of maize grain yield from near infrared reflectance spectroscopy of kernels. Plant Phenome Journal, 3(1). 10.1002/ppj2.20002

Li, Z., Taylor, J., Yang, H., Casa, R., Jin, X., Li, Z., Song, X., & Yang, G. (2020). A hierarchical interannual wheat yield and grain protein prediction model using spectral vegetative indices and meteorological data. Field Crops Research, 248(107711), 107711.

Magney, T. S., Brissette, L. E. G., Pierrat, Z. A., Logan, B., Reblin, J., Nelson, S., Stutz, J., Frankenberg, C., Bowling, D. R., & Wong, C. Y. S. (2026). Tracking subtle seasonal shifts in pigment composition with hyperspectral reflectance in a temperate evergreen forest. Tree Physiology, 46(1). 10.1093/treephys/tpaf108

Manetas, Y. (2006). Why some leaves are anthocyanic and why most anthocyanic leaves are red? Flora, 201(3), 163–177.

Mantel, N. (1967). The detection of disease clustering and a generalized regression approach. Cancer Research, 27(2_Part_1), 209–220.

Mbebi, A. J., Breitler, J.-C., Bordeaux, M., Sulpice, R., McHale, M., Tong, H., Toniutti, L., Castillo, J. A., Bertrand, B., & Nikoloski, Z. (2022). A comparative analysis of genomic and phenomic predictions of growth-related traits in 3-way coffee hybrids. G3 (Bethesda, Md.), 12(9). 10.1093/g3journal/jkac170

Montesinos-López, A., Montesinos-López, O. A., Montesinos-López, J. C., Flores-Cortes, C. A., de la Rosa, R., & Crossa, J. (2021a). A guide for kernel generalized regression methods for genomic-enabled prediction. Heredity, 126(4), 577–596.

Montesinos-López, A., Montesinos-López, O. A., Montesinos-López, J. C., Flores-Cortes, C. A., de la Rosa, R., & Crossa, J. (2021b). A guide for kernel generalized regression methods for genomic-enabled prediction. Heredity, 126(4), 577–596.

Morota, G., & Gianola, D. (2014). Kernel-based whole-genome prediction of complex traits: a review. Frontiers in Genetics, 5, 363.

Oliveira, L. F. R. de, & Santana, R. C. (2020). Exploratory analysis of nutrient concentrations in Eucalyptus leaf color patterns. Advances in Forestry Science, 7(2), 973–979.

Paulus, S., & Mahlein, A.-K. (2020). Technical workflows for hyperspectral plant image assessment and processing on the greenhouse and laboratory scale. GigaScience, 9(8). 10.1093/gigascience/giaa090

Pérez, P., & de los Campos, G. (2014). Genome-wide regression and prediction with the BGLR statistical package. Genetics, 198(2), 483–495.

Poland, J. A., Brown, P. J., Sorrells, M. E., & Jannink, J.-L. (2012). Development of high-density genetic maps for barley and wheat using a novel two-enzyme genotyping-by-sequencing approach. PloS One, 7(2), e32253.

Riedelsheimer, C., Lisec, J., Czedik-Eysenberg, A., Sulpice, R., Flis, A., Grieder, C., Altmann, T., Stitt, M., Willmitzer, L., & Melchinger, A. E. (2012). Genome-wide association mapping of leaf metabolic profiles for dissecting complex traits in maize. Proceedings of the National Academy of Sciences of the United States of America, 109(23), 8872–8877.

Rife, T. W., Courtney, C., Hershberger, J., Gore, M. A., Neilsen, M., & Poland, J. (2021). Prospector: A mobile application for portable, high throughput near infrared spectroscopy phenotyping. The Plant Phenome Journal, 4(1). 10.1002/ppj2.20024

Rincent, R., Charpentier, J.-P., Faivre-Rampant, P., Paux, E., Le Gouis, J., Bastien, C., & Segura, V. (2018a). Phenomic Selection Is a Low-Cost and High-Throughput Method Based on Indirect Predictions: Proof of Concept on Wheat and Poplar. G3 (Bethesda, Md.), 8(12), 3961–3972.

Rincent, R., Charpentier, J.-P., Faivre-Rampant, P., Paux, E., Le Gouis, J., Bastien, C., & Segura, V. (2018b). Phenomic Selection Is a Low-Cost and High-Throughput Method Based on Indirect Predictions: Proof of Concept on Wheat and Poplar. G3 (Bethesda, Md.), 8(12), 3961–3972.

Rincent, R., Charpentier, J.-P., Faivre-Rampant, P., Paux, E., Le Gouis, J., Bastien, C., & Segura, V. (2018c). Phenomic Selection Is a Low-Cost and High-Throughput Method Based on Indirect Predictions: Proof of Concept on Wheat and Poplar. G3 (Bethesda, Md.), 8(12), 3961–3972.

Rincent, R., Solin, J., Lorenzi, A., Nunes, L., Griveau, Y., Pirus, L., Kermarrec, D., Bauland, C., Reymond, M., & Moreau, L. (2025). Using phenomic selection to predict hybrid values with NIR spectra measured on the parental lines: proof of concept on maize. TAG. Theoretical and Applied Genetics. Theoretische Und Angewandte Genetik, 138(1), 28.

Roscher-Ehrig, L., Weber, S. E., Abbadi, A., Malenica, M., Abel, S., Hemker, R., Snowdon, R. J., Wittkop, B., & Stahl, A. (2024). Phenomic Selection for Hybrid Rapeseed Breeding. Plant Phenomics (Washington, D.C.), 6, 0215.

Rowe, C. L. J., & Leger, E. A. (2011). Competitive seedlings and inherited traits: a test of rapid evolution of Elymus multisetus (big squirreltail) in response to cheatgrass invasion. Evolutionary Applications, 4(3), 485–498.

Rusch, H. L., Hunter, M. C., Kraus, A., Tautges, N. E., & Jungers, J. M. (2025). Intermediate wheatgrass as a dual use crop for grain and grazing. Frontiers in Agronomy, 7(1534962). 10.3389/fagro.2025.1534962

Rweyongeza, D. M., Yeh, F. C., & Dhir, N. K. (2004). Genetic parameters for seasonal height and height growth curves of white spruce seedlings and their implications to early selection. Forest Ecology and Management, 187(2-3), 159–172.

Sakiroglu, M., Dong, C., Hall, M. B., Jungers, J., & Picasso, V. (2020). How does nitrogen and forage harvest affect belowground biomass and nonstructural carbohydrates in dual use Kernza intermediate wheatgrass? Crop Science, 60(5), 2562–2573.

Sarić, R., Nguyen, V. D., Burge, T., Berkowitz, O., Trtílek, M., Whelan, J., Lewsey, M. G., & Čustović, E. (2022). Applications of hyperspectral imaging in plant phenotyping. Trends in Plant Science, 27(3), 301–315.

Schuhl, H., Brown, K. E., Sheng, H., Bhatt, P. K., Gutierrez, J., Schneider, D., Casto, A. L., Acosta-Gamboa, L., Ballenger, J. G., Barbero, F., Braley, J., Brown, A. M., Chavez, L., Cunningham, S., Dilhara, M., Dimech, A. M., Duenwald, J. G., Fischer, A., Gordon, J. M., … Fahlgren, N. (2025). PlantCV v4: Image analysis software for high-throughput plant phenotyping. In bioRxiv. bioRxiv. 10.1101/2025.11.19.689271

Seyum, E. G., Bille, N. H., Abtew, W. G., Munyengwa, N., Bell, J. M., & Cros, D. (2022). Genomic selection in tropical perennial crops and plantation trees: a review. Molecular Breeding : New Strategies in Plant Improvement, 42(10), 58.

Sharda, S., Kumar, S., Setia, R., Dhiman, P., Patel, N. R., Pateriya, B., Salem, A., & Elbeltagi, A. (2025). Evaluation of different spectral indices for wheat lodging assessment using machine learning algorithms. Scientific Reports, 15(1), 21774.

Shi, T., Zhu, A., Jia, J., Hu, X., Chen, J., Liu, W., Ren, X., Sun, D., Fernie, A. R., Cui, F., & Chen, W. (2020). Metabolomics analysis and metabolite-agronomic trait associations using kernels of wheat (Triticum aestivum) recombinant inbred lines. The Plant Journal : For Cell and Molecular Biology, 103(1), 279–292.

Silva-Perez, V., Molero, G., Serbin, S. P., Condon, A. G., Reynolds, M. P., Furbank, R. T., & Evans, J. R. (2018). Hyperspectral reflectance as a tool to measure biochemical and physiological traits in wheat. Journal of Experimental Botany, 69(3), 483–496.

Sleper, J. A., Zheng, C., Ji, L., Wang, X., Abd-Elrahman, A., & Whitaker, V. M. (2025). Exploring the efficacy of phenomic and genomic selection for yield and fruit quality traits in strawberry. The Plant Genome, 18(4), e70156.

Smaje, C. (2015). The Strong Perennial Vision: A Critical Review. Agroecology and Sustainable Food Systems, 39(5), 471–499.

Sthapit, S. R., Crain, J., Larson, S., Anderson, J. A., Bajgain, P., DeHaan, L. R., & Poland, J. (2025). A low-coverage skim-sequencing and imputation pipeline for genomic selection. The Plant Genome, 18(4), e70139.

Sun, Y., Tong, C., He, S., Wang, K., & Chen, L. (2018). Identification of nitrogen, phosphorus, and potassium deficiencies based on temporal dynamics of leaf morphology and color. Sustainability, 10(3), 762.

Tuberosa, R. (2012). Phenotyping for drought tolerance of crops in the genomics era. Frontiers in Physiology, 3, 347.

Umaña, M. N., Needham, J., & Fortunel, C. (2025). From seedlings to adults: Linking survival and leaf functional traits over ontogeny. Ecology, 106(1), e4469.

Van der Laan, L., Parmley, K., Saadati, M., Pacin, H. T., Panthulugiri, S., Sarkar, S., Ganapathysubramanian, B., Lorenz, A., & Singh, A. K. (2025). Genomic and phenomic prediction for soybean seed yield, protein, and oil. The Plant Genome, 18(1), e70002.

Van Tassel, D. L., DeHaan, L. R., & Cox, T. S. (2010). Missing domesticated plant forms: can artificial selection fill the gap? Evolutionary Applications, 3(5-6), 434–452.

Villa, P., Bolpagni, R., Pinardi, M., & Tóth, V. R. (2021). Leaf reflectance can surrogate foliar economics better than physiological traits across macrophyte species. Plant Methods, 17(1), 115.

Wagoner, P. (1990). Perennial Grain. Journal of Soil and Water Conservation, 45(1), 81–82.

Wagoner, P., & Schaeffer, J. R., Dr. agr. (1990). Perennial grain development: Past efforts and potential for the future. Critical Reviews in Plant Sciences, 9(5), 381–408.

Wang, F., Feldmann, M. J., & Runcie, D. E. (2025). Do not benchmark phenomic prediction against genomic prediction accuracy. Plant Phenome Journal, 8(1). 10.1002/ppj2.70029

Wang, X., Xing, E. P., & Schaid, D. J. (2015). Kernel methods for large-scale genomic data analysis. Briefings in Bioinformatics, 16(2), 183–192.

Werling, B. P., Dickson, T. L., Isaacs, R., Gaines, H., Gratton, C., Gross, K. L., Liere, H., Malmstrom, C. M., Meehan, T. D., Ruan, L., Robertson, B. A., Robertson, G. P., Schmidt, T. M., Schrotenboer, A. C., Teal, T. K., Wilson, J. K., & Landis, D. A. (2014). Perennial grasslands enhance biodiversity and multiple ecosystem services in bioenergy landscapes. Proceedings of the National Academy of Sciences of the United States of America, 111(4), 1652–1657.

Winn, Z. J., Amsberry, A. L., Haley, S. D., DeWitt, N. D., & Mason, R. E. (2023). Phenomic versus genomic prediction—A comparison of prediction accuracies for grain yield in hard winter wheat lines. Plant Phenome Journal, 6(1). 10.1002/ppj2.20084

Woeltjen, S., Hanlon, M., Brown, K., Schuhl, H., Baxter, I., & Miller, A. (2026). Smartphone image capture system and image analysis pipelines enable accurate and efficient phenotyping of spaced plant mapping populations. In bioRxiv. bioRxiv. 10.64898/2026.01.06.697986

Xu, L., Chen, L., Luo, Q., Zhao, S., Huang, J., Wang, K., Yang, Z., Weng, X., Fang, K., & Feng, H. (2025). Leveraging UAV hyperspectral imaging for crop physiology and biochemistry: A comprehensive review of feature extraction and selection methods. Plant Phenomics (Washington, D.C.), 100141, 100141.

Xu, R., Ferguson, J., Breil-Aubert, M., Kromdijk, J., & Nikoloski, Z. (2026). Generalizability and transferability of machine learning models using hyperspectral reflectance data for maize traits. Scientific Reports, 16(1), 5865.

Yang, W., Wang, S., Zhao, X., Zhang, J., & Feng, J. (2015). Greenness identification based on HSV decision tree. Information Processing in Agriculture, 2(3-4), 149–160.

Yendrek, C. R., Tomaz, T., Montes, C. M., Cao, Y., Morse, A. M., Brown, P. J., McIntyre, L. M., Leakey, A. D. B., & Ainsworth, E. A. (2017). High-Throughput Phenotyping of Maize Leaf Physiological and Biochemical Traits Using Hyperspectral Reflectance. Plant Physiology, 173(1), 614–626.

Zargar, S. M., Manzoor, M., Bhat, B., Wani, A. B., Sofi, P. A., Sudan, J., Ebinezer, L. B., Dall’Acqua, S., Peron, G., & Masi, A. (2023). Metabolic-GWAS provides insights into genetic architecture of seed metabolome in buckwheat. BMC Plant Biology, 23(1), 373.

Zhang, H., Ge, Y., Xie, X., Atefi, A., Wijewardane, N. K., & Thapa, S. (2022). High throughput analysis of leaf chlorophyll content in sorghum using RGB, hyperspectral, and fluorescence imaging and sensor fusion. Plant Methods, 18(1), 60.

Zhang, S., Huang, G., Zhang, Y., Lv, X., Wan, K., Liang, J., Feng, Y., Dao, J., Wu, S., Zhang, L., Yang, X., Lian, X., Huang, L., Shao, L., Zhang, J., Qin, S., Tao, D., Crews, T. E., Sacks, E. J., … Hu, F. (2022). Sustained productivity and agronomic potential of perennial rice. Nature Sustainability, 6(1), 28–38.

Zhang, X., Sallam, A., Gao, L., Kantarski, T., Poland, J., DeHaan, L. R., Wyse, D. L., & Anderson, J. A. (2016). Establishment and Optimization of Genomic Selection to Accelerate the Domestication and Improvement of Intermediate Wheatgrass. The Plant Genome, 9(1). 10.3835/plantgenome2015.07.0059

